# Attentional modulation is orthogonal to disinhibition by VIP interneurons in primary visual cortex

**DOI:** 10.1101/2022.11.28.518253

**Authors:** Dylan Myers-Joseph, Katharina A. Wilmes, Marian Fernandez-Otero, Claudia Clopath, Adil G. Khan

## Abstract

Attentional modulation of sensory processing is a key feature of cognition, yet its neural circuit basis is poorly understood. A candidate mechanism is the disinhibition of pyramidal cells through vasoactive intestinal peptide (VIP) and somatostatin (SOM) positive interneurons. However, the interaction of attentional modulation and VIP-SOM disinhibition has never been directly tested. We used all-optical methods to bi-directionally manipulate VIP interneuron activity as mice performed an attention switching task. We measured the activity of VIP, SOM and parvalbumin (PV) positive interneurons and pyramidal neurons identified in the same tissue and found that although activity in all cell classes was modulated by both attention and VIP manipulation, their effects were orthogonal. Attention and VIP-SOM disinhibition relied on distinct patterns of changes in activity and reorganisation of interactions between inhibitory and excitatory cells. Circuit modelling revealed a precise network architecture consistent with multiplexing strong yet non-interacting modulations in the same neural population.

## Introduction

Attention exerts a powerful influence on how cortical circuits process sensory information. There are multiple forms of visual attention including spatial, feature based, and cross modality, and in all these cases attention can modulate visual processing leading to improved behavioural performance^1–5^. Typically, stimulus selectivity in visual cortex increases with attention, and this increased selectivity is believed to underlie the improved behavioural discriminability of attended stimuli^2,5–8^. The increase in stimulus selectivity of neurons occurs prominently through changes in stimulus-evoked firing rates. These changes can be increases^7–9^ or decreases^10,11^ in evoked firing rates or combinations of these^5^, which ultimately result in more selective stimulus representations. Changes in noise correlations, variability and synchrony between neurons may also play a role in improving the discriminability of stimuli by downstream circuits^12–14^. However, despite this thorough phenomenological description of the ways in which attention influences visual processing, very little is known about the circuit basis of attentional modulation in visual cortex.

Local inhibitory interneurons play a crucial role in shaping cortical activity. Cortical activity is processed by interconnected networks of cells containing multiple classes of excitatory and GABAergic inhibitory interneurons with distinct molecular, cellular and connectivity properties^15–19^. VIP interneurons are key regulators of cortical function, most prominently due to their strong inhibition of SOM interneurons which results in disinhibition of excitatory pyramidal cells^17,20,21^. VIP-SOM driven disinhibition provides useful computational opportunities^22,23^. On a longer timescale, VIP-SOM disinhibition gates the plasticity of inputs onto excitatory cells^24–27^. Better studied, however, is the immediate effect of increasing activity in VIP interneurons, which leads to enhanced firing of local pyramidal cells^23,28^. This effect has been observed across many regions of sensory^20,21,29–34^, motor^35^ and prefrontal cortex^36,37^.

Disinhibitory modulation of sensory cortex through VIP and SOM interneurons has in fact been proposed to be the core mechanism underlying attentional modulation in visual cortex^38,39^. In this account, projections from frontal cortex activate VIP interneurons in V1 which leads to enhanced stimulus-evoked activation of pyramidal (PYR) cells though VIP-SOM disinhibition, and may lead to subsequent improvement in visual behaviour^40^. Attentional modulation in V1 and visual discrimination behaviour may also depend on cholinergic inputs to visual cortex^41–43^, and cortical VIP interneurons are activated by acetylcholine^44,45^, raising the possibility of VIP interneurons being a key player in cholinergic visual gain enhancement during attention^46^. This account has parallels with the suggestion that locomotion elevates the gain of visual responses in V1 through VIP-SOM disinhibition^1,29,47^. However, the direct interaction of attentional modulation and VIP modulation has never been tested.

If attention acts through VIP-SOM disinhibition, then two broad predictions exist. First, attention should have similar effects on pyramidal neurons as manipulating VIP interneurons. As a consequence, the degree to which attention modulates stimulus selectivity should depend on whether the VIP-SOM disinhibitory circuit is activated or not. In other words, there should be an interaction between attentional modulation and VIP-SOM disinhibition. Second, there should be similarities in how attention and VIP-SOM disinhibition modify the activity of different cell classes and the interactions between them.

To test these predictions, we used an all-optical approach in V1, where we simultaneously imaged the activity of PYR, PV, SOM and VIP interneurons, while optogenetically manipulating VIP interneurons to mimic physiologically relevant levels of activity, as mice performed a cross-modal attention-switching task. As mice switched between attending to or ignoring the same pair of visual stimuli, we observed robust attentional modulation of stimulus selectivity in V1. We also observed strong enhancement of V1 activity when photoactivating VIP interneurons, confirmed to be through VIP-SOM disinhibition, and a reduction in V1 activity when photoinhibiting VIP interneurons. However, when we combined VIP manipulations with attentional changes, we found that the two modulations did not interact, and instead were orthogonal. The changes induced by these two modulations occurred through distinct mechanisms: while VIP photoactivation led to predominantly enhanced activity in VIP, PV and PYR neurons, and largely suppressed activity in SOM neurons, attention led to heterogeneous changes, marked by a prominent reduction in evoked activity in all four cell classes. Circuit modelling revealed that only a specific network architecture can account for the experimental findings. These results demonstrate that VIP-SOM disinhibition does not underlie attentional modulation of stimulus selectivity in V1. At the same time, they reveal a remarkably versatile cortical circuit, which allows multiplexing of multiple non-interacting signals on the same neural populations.

## Results

### Modulation of stimulus selectivity by attention in V1

To study the neural circuit basis of attentional modulation, we trained mice to perform a cross-modality attention switching task (Fig. 1a, b). Head-fixed mice switched between blocks of visual discrimination and olfactory discrimination in which they licked a reward spout to obtain a reward in response to one of two visual grating stimuli or odours. During the olfactory discrimination blocks, the same grating stimuli used in the visual discrimination blocks were presented on 70% of trials but were irrelevant to the task (Fig. 1b). Mice attended to and accurately discriminated the grating stimuli in the visual block but ignored the same grating stimuli while successfully discriminating odours during the olfactory blocks (Fig. 1c, behavioural d’ of visual discrimination: visual block, attend visual, 3.47 ± 0.77; olfactory block, ignore visual, 0.55 ± 0.87, Wilcoxon signed-rank test p = 7.56×10^−10^; behavioural d’ of olfactory discrimination, 3.90 ± 0.67, n = 50 sessions).

**Figure 1:**
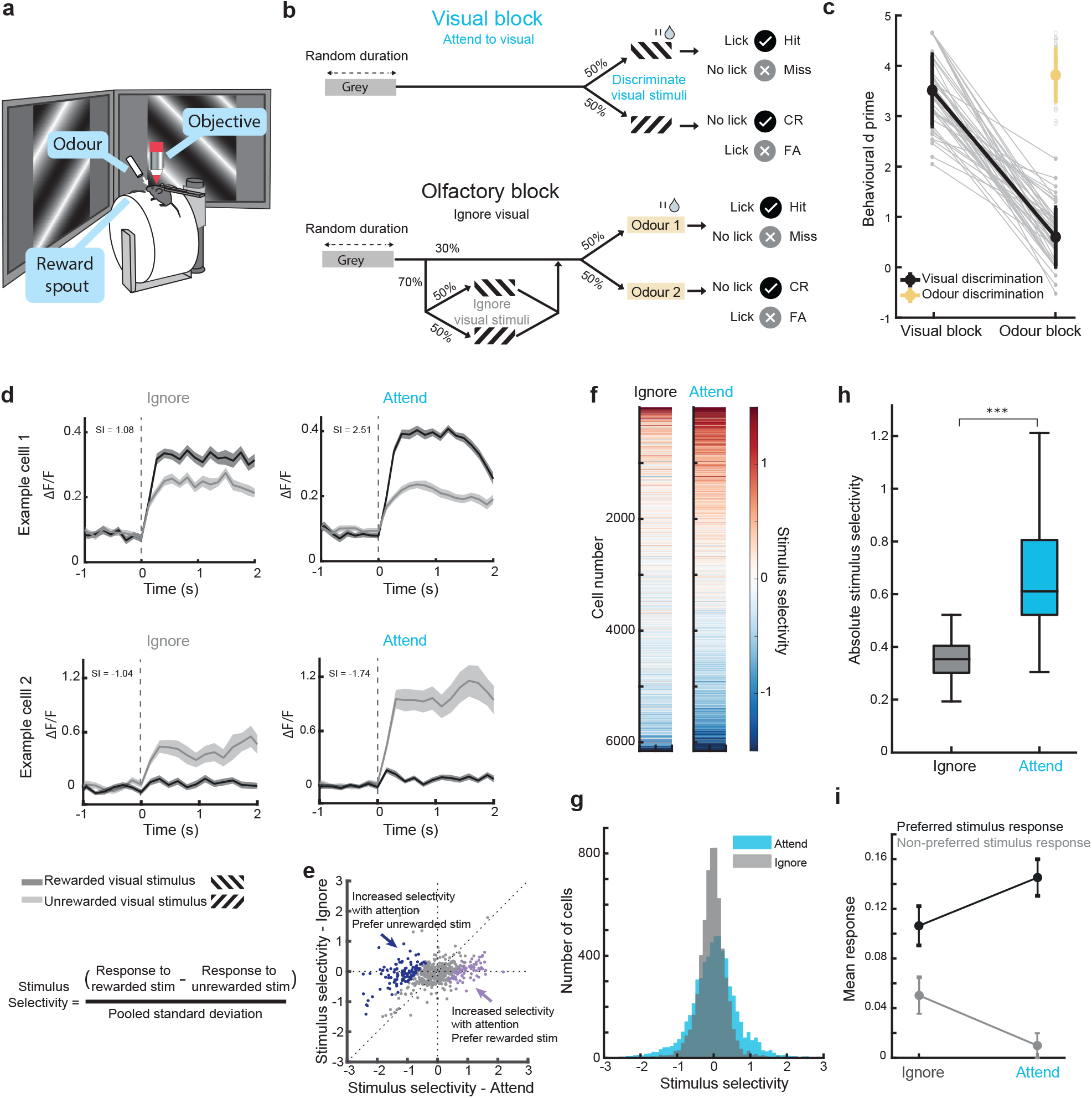
Modulation of stimulus selectivity in V1 neurons during an attention-switching task **a)** Schematic of the experimental apparatus. **b**) Schematic of the behavioural task. Top, visual block: mice were rewarded for licking the reward spout when gratings of a specific orientation were presented (+15 degrees from vertical, rewarded grating) and not when gratings of a second orientation were presented (−15 degrees from vertical, unrewarded grating). Olfactory block: mice were rewarded for licking when odour 1 was presented and not when odour 2 or either visual grating was presented. **c**) Behavioural discrimination performance (behavioural d’) across attention (n = 50 sessions, 15 mice, Wilcoxon signed-rank test between attended visual trials and ignored visual trials, p = 7.55×10^−10^). Connected points indicate visual discrimination, individual points in the odour block represent olfactory discrimination. Grey lines and points are individual sessions, coloured lines show the average of all sessions, error bars indicate STD. **d**) Average responses from 2 example cells to the rewarded and unrewarded visual gratings in the odour and visual blocks, showing an increase in selectivity in the attend condition. Top, a cell with a preference for the rewarded grating stimulus (positive selectivity). Bottom, a cell with a preference for the unrewarded grating stimulus (negative selectivity). Numbers indicate stimulus selectivity. **e**) Stimulus selectivity of all non-VIP cells (calculated with stimulus evoked responses averaged 0-1s from stimulus onset) when attending or ignoring the same stimuli from an example session. Negatively selective cells that significantly increase their selectivity with attention are highlighted in dark blue. Positively selective cells that significantly increase their selectivity with attention are highlighted in purple. **f**) Stimulus selectivity of the same cells in the attend and ignore conditions (columns). Cells were ordered by their mean selectivity across both contexts (n = 6153 neurons, 15 mice). **g**) Histograms of stimulus selectivity when ignoring and attending the visual stimuli (n = 6153 neurons, 15 mice). **h**) Box plots of absolute stimulus selectivity during the ignore and attend conditions (n = 50 sessions, 15 mice, p = 7.56×10^−10^, Wilcoxon signed-rank test). **i**) Average baseline subtracted responses to the preferred and non-preferred visual stimuli in the ignore and attend conditions for cells that significantly increased their stimulus selectivity with attention (n = 50 sessions, 15 mice, error bars indicate SEM).

We expressed the calcium indicator GCaMP7f^48^ in V1 using adeno-associated virus (AAV) vectors and measured responses of layer 2/3 neurons using two-photon calcium imaging during the task. We compared the responses to the same pair of visual stimuli in the attend and ignore conditions, and observed a robust modulation of stimulus responses with attention (Fig. 1d). These response changes modified stimulus selectivity (difference in the responses to the rewarded and unrewarded grating stimuli, normalized by the pooled standard deviation) such that cells preferring both the rewarded and unrewarded grating stimuli showed increased stimulus selectivity (Fig. 1d, e). Across the population, we observed individual cells increasing their stimulus selectivity (Fig. 1f) and a broadening of the distribution of stimulus selectivity with attention due to more positive and more negative values, indicating higher preference for the rewarded and non-rewarded stimuli respectively (Fig. 1g). Consequently, the absolute stimulus selectivity of the neural population increased significantly with attention, as reported previously^5,49^ (Fig. 1h, average absolute selectivity ignore, 0.35 ± 0.10 median ± IQR, attend 0.61 ± 0.28, Wilcoxon signed-rank test, P = 7.56×10^−10^, N = 50 sessions, 15 mice). The increase in stimulus selectivity could not be accounted for by changes in running or licking behaviour, and remained after removing cells most influenced by running and licking (Supplementary Fig. 1). We restricted the analysis to cells which significantly increased their selectivity for either stimulus with attention and found that a combination of increased responses to the preferred and decreased responses to the non-preferred stimuli led to the increase in average selectivity (Fig. 1i).

These results established that visual stimulus selectivity in V1 was strongly modulated in this attention switching task. A candidate mechanism for this selectivity modulation is VIP interneuron mediated disinhibition^38^. To test the role of VIP interneurons in attentional modulation, we first asked whether optogenetic VIP activation within physiological levels modified neural activity in the same mice passively viewing stimuli.

### VIP activation strongly modulates cortical responses

To study the effect of activating VIP interneurons on visual stimulus responses, we expressed the red-shifted excitatory opsin Chrimson selectively in VIP interneurons and GCaMP7f non-selectively in V1 neurons using AAVs (Fig. 2a). We used an all-optical approach where we photoactivated VIP interneurons while measuring the activity of VIP and non-VIP cells in the same local circuit (Fig. 2a, the population of all non-VIP cells is dominated by pyramidal neurons^18^, see also Fig. 7 for analysis of identified cell classes)

**Figure 2:**
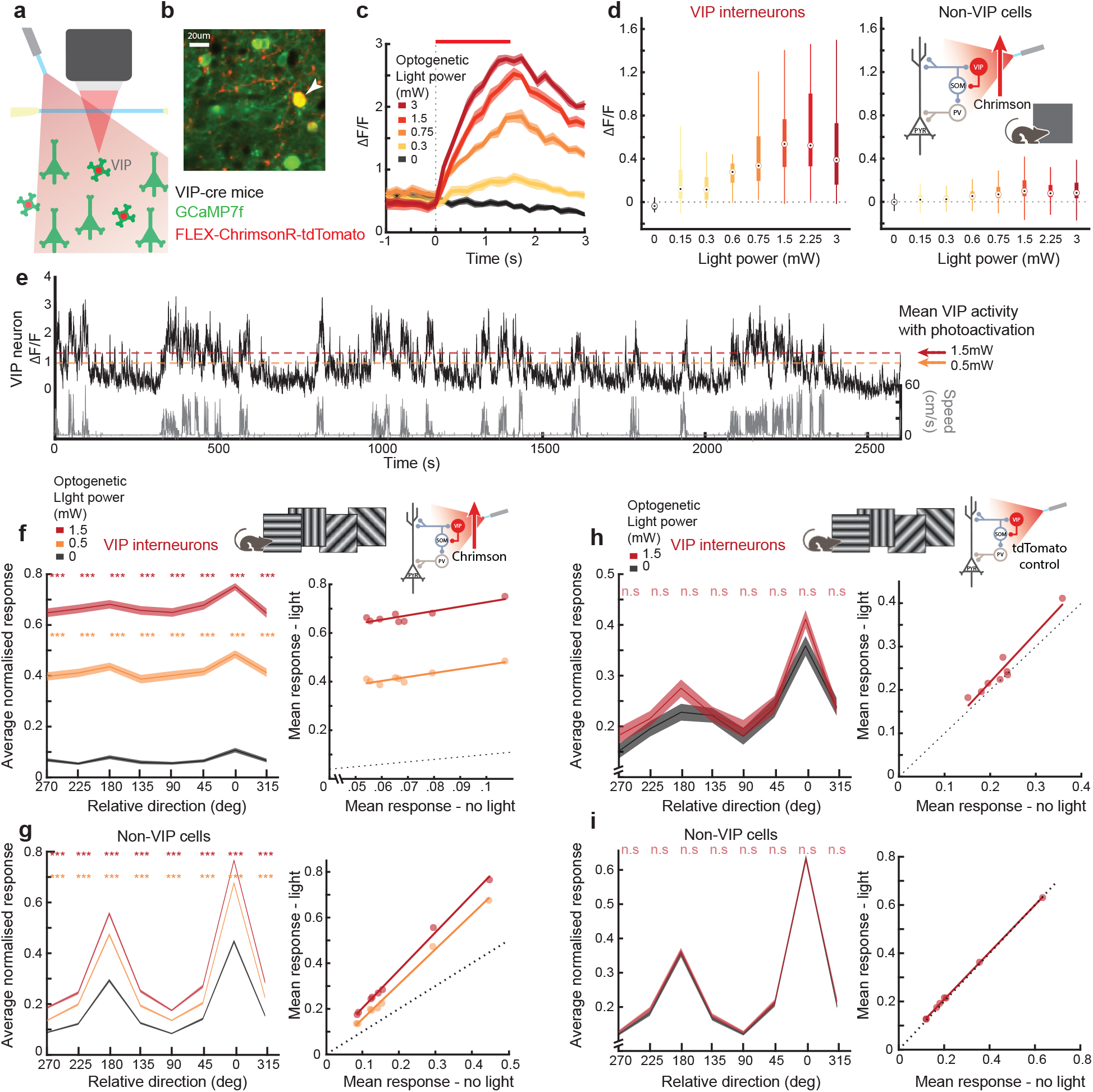
VIP activation leads to multiplicative increase in activity of non-VIP cells **a**) Schematic of near-simultaneous imaging and optogenetic stimulation. **b**) Example region of an *in vivo* imaged plane showing all neurons expressing GCaMP7f and a VIP interneuron (arrowhead) additionally expressing Chrimson-tdTomato. **c**) Mean responses of an example VIP interneuron to different light powers. Responses are aligned to optogenetic light onset (dashed line). Red bar indicates optogenetic stimulation duration (1.5s), shading indicates SEM. **d**) Box plots of optogenetically evoked activity (mean 0-1s, baseline subtracted) across different light powers for VIP interneurons (left, n = 91 cells, 6 mice) and non-VIP cells (right, n = 2233 cells, 6 mice). Inset: schematic of the VIP-SOM disinhibitory circuit (studied further below) with VIP interneuron activation during passive grey screen viewing. **e**) Example activity trace showing running evoked activity of an example VIP interneuron (black) and running speed (grey), dashed lines indicate mean optogenetically evoked activity in VIP interneurons (orange, low power, red, high power). **f**) Left, stimulus evoked normalised activity in response to different oriented drifting gratings, averaged across all VIP interneurons (n = 141 cells) aligned to their preferred direction. ***, p<0.001 Wilcoxon signed-rank test for photoactivation compared to non-photoactivation conditions at each direction, corrected for multiple comparisons. Right, the same data shown as average activity with and without optogenetic light. Linear regression, low power, slope = 1.823, intercept = 0.327. High power, slope = 2.081, intercept = 0.52, 8 mice. **g**) Same as f, for all orientation selective non-VIP neurons, n = 1044 cells. Low power, slope = 1.568, intercept = 0.009. High power, slope = 1.634, intercept = 0.032, 8 mice. **h, i**) Same as f, g, for control mice expressing only tdTomato and no opsin, n.s. indicates non-significant, n = 61 VIP interneurons, and 434 non-VIP cells, 3 mice.

VIP interneurons were robustly activated by increasing light powers *in vivo* (Fig. 2c). For each imaged site, we conducted a calibration session in which we increased the light power until we reached a plateau of VIP interneuron activation (Fig. 2d) and chose a ‘low’ and ‘high’ light power based on the shape of this curve (see methods). These light powers were typically 0.6 mW (low, range 0.5 to 0.6 mW) and 1.5 mW (high, range 1.5 to 2.25 mW) although tailored values were selected for each site. We established that our photoactivation was within a physiological range by comparing the response amplitude of VIP interneurons during photoactivation and during spontaneous running bouts. Both low and high light powers evoked activity in VIP interneurons which spanned the activity range naturally observed during locomotion (Fig. 2e, mean VIP interneuron activity with low and high light power, 1.32 and 1.77 ΔF/F respectively. Mean 75^th^ and 95^th^ percentiles of activity during locomotion without photostimulation for all VIP interneurons, 1.23, and 2.06 ΔF/F respectively).

We photoactivated VIP interneurons while we passively presented a range of oriented drifting grating stimuli to the mice. We confirmed that VIP interneurons increased their activity at each grating direction with increasing light power (Fig. 2f). As a result, non-VIP cell stimulus responses were strongly positively modulated at each direction (Fig. 2g). By comparing the average responses at each direction, we found that VIP interneuron activation caused a largely multiplicative enhancement in orientation-tuned non-VIP cells (Fig. 2g right, linear regression here and below, low power: slope = 1.568, intercept = 0.009, high power: slope = 1.634, intercept = 0.032, n = 1044 cells, 8 mice). Identical analysis on control mice with the same light stimulation but no opsin expression did not show any significant changes in activity, ruling out light-evoked artefacts at this light power (Fig. 2 h, i). Since a purely multiplicative increase in activity is unlikely to lead to changes in stimulus selectivity ^9^, these results indicated that VIP interneurons may not be involved in the stimulus selectivity modulations we observed with attention. However, directly testing this hypothesis required studying VIP activity during the attention-switching task.

### Testing the interaction between VIP activation and attention

To understand the role of VIP interneurons in attentional modulation, we first asked to what extent the activity of VIP interneurons themselves was modulated in this attention switching task. We compared the attend and ignore conditions and found a small but significant difference in pre-stimulus VIP interneuron activity, with higher activity in the attend condition (Supplementary Fig. 2a). However, this could be accounted for by a difference in pre-stimulus running speed in the two blocks (Supplementary Fig. 2b), and correcting for the pre-stimulus activity resulted in no significant difference in stimulus evoked VIP interneuron activity between the attend and ignore conditions (Supplementary Fig. 2c). This suggested that VIP interneurons are not important in producing attentional modulation.

However, to conclusively determine whether VIP activity underlies attentional modulation, we photoactivated VIP interneurons in randomly interleaved trials while mice performed the attention-switching task with the light-on period from 100ms before the grating stimulus onset until 1.5s after grating stimulus onset (Fig. 3a). Photoactivation had no effect on the behaviour itself (Supplementary Fig. 3a, b) possibly because the contralateral hemisphere was unperturbed. The activity of VIP interneurons during the task was again enhanced with increasing light power, both when the visual stimuli were attended and ignored (Fig. 3b, top), and this led to an increase in non-VIP cell stimulus evoked activity (Fig. 3b, bottom).

**Figure 3.**
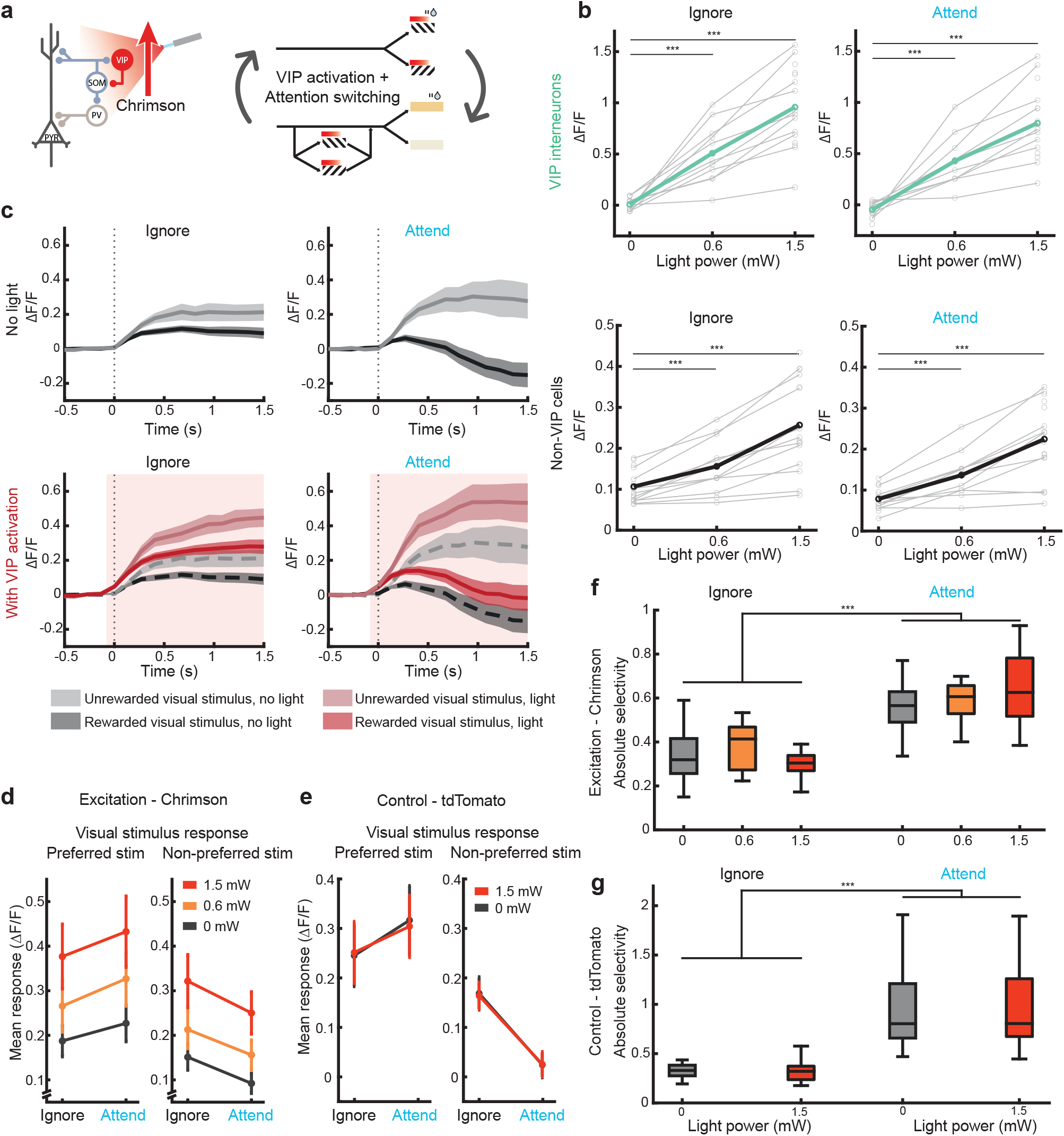
No interaction between VIP modulation and attentional modulation. **a**) Schematic showing VIP photoactivation during the attention switching task. Light onset (red bars) was from -0.1s to 1.5s relative to visual stimulus onset. Light was ramped off over 0.2s. **b**) Mean visual stimulus evoked activity with increasing VIP photoactivation. Top, all VIP interneurons, bottom, all non-VIP cells. Left, responses when ignoring the visual stimuli, right, responses when attending the visual stimuli. Wilcoxon signed-rank test between photoactivation and non-photoactivation conditions, ***, p< 0.001, n = 17 sessions, 7 mice. Gray lines indicate individual session averages, coloured lines indicate overall average. **c**) Top, mean visual stimulus evoked activity for all non-VIP cells with preference for the unrewarded stimulus that significantly increased their selectivity with attention (mean of n=17 session averages, shading indicates SEM). Bottom, same sessions, responses with additional VIP photoactivation (red). Responses from top are superimposed for comparison (grey dashed lines, light red shading indicates light onset). **d**) Black, mean visual stimulus evoked activity (averaged 0-1s) of all non-VIP cells with preference for the unrewarded stimulus that significantly increased their selectivity with attention, n = 17 sessions. Orange and red, same responses with additional VIP photoactivation. Error bars indicate SEM. **e**) Same as d for control mice expressing tdTomato, n = 17 sessions, 3 mice. **f**) Absolute stimulus selectivity with increasing VIP photoactivation for all non-VIP cells which significantly increased their selectivity with attention (n = 17 sessions). Stimulus selectivity measured when ignoring the visual stimuli (left) and attending the same stimuli (right). There was a significant effect of attention on selectivity, but not of VIP activation or an interaction between the two (2-way ANOVA, attention p = 2.44×10^−11^, other ps > 0.05). **g**) Same as f for control mice expressing tdTomato, n = 17 sessions, 3 mice.

To study the interaction of VIP activation with attentional modulation of stimulus selectivity, we selected all non-VIP neurons which showed a significant increase in stimulus selectivity with attention. The responses of this population to the two grating stimuli in the ignore and attend conditions, with and without VIP photoactivation showed modulatory effects of both attention and VIP activation (Fig. 3c, cells with preference for the unrewarded stimulus, similar effects seen for cells with a preference for the rewarded stimulus, data now shown). Increasing light power evenly increased average (0 to 1s) activity of this population in both ignore and attend conditions, for both the preferred and unpreferred stimulus (Fig. 3d). We performed a two-way ANOVA and found a significant main effect for both attention and VIP activation, but no significant interaction effect (rewarded stimulus: main effect for attention p = 1.31×10^−05^ and VIP activation p = 1.70×10^−05^, no interaction effect p = 0.688). Similar results of no interaction between attention and VIP activation were obtained for the unrewarded stimulus and from cells with a preference for the rewarded stimulus (data not shown), as well as in control mice (Fig. 3e, see also Supplementary Tables 1-6).

Having found no interaction between attention and VIP activation on stimulus evoked responses, we asked whether there was any change in stimulus selectivity with VIP activation in either the ignore or attend condition (Fig. 3f). We performed a two-way ANOVA on the data in Fig. 3f (all non-VIP cells which significantly increased their selectivity with attention) and found a significant main effect on selectivity from attention (p = 2.44×10^−11^), but not from VIP activation (p = 0.46) and no interaction effect (p = 0.15). The same result was obtained when taking all non-VIP cells (a significant main effect on selectivity from attention, p = 4.50×10^−4^, but not from VIP activation, p = 0.06 and no interaction effect, p = 0.33). Crucially, in the control mice expressing tdTomato only (Fig. 3g), there was also only a significant effect on selectivity from attention (p = 2.69×10^−17^), but not from light delivery (p = 0.36) and no interaction effect (p = 0.48). Thus, attention and VIP-driven disinhibition both induce robust modulation of the same neural population, but the two effects do not interact.

### VIP inhibition leads to a modest suppression of cortical responses during passive viewing

Although the above results demonstrated that VIP activation did not interact with attentional modulation of stimulus selectivity, to better understand the role of VIP interneurons in shaping cortical stimulus evoked responses we next studied the effect of inhibiting VIP interneurons. We used a similar all-optical approach, by expressing the inhibitory opsin ArchT in VIP interneurons (Fig. 4a). VIP interneurons were progressively inhibited by increasing light powers *in vivo* (Fig. 4b). As with VIP activation, we conducted a calibration session for each site in which we increased the light power until we reached a floor of VIP inhibition (Fig. 4c) and chose a single light power based on the shape of this curve, typically 1.5mW (range 1.15 to 1.8mW, see methods).

**Figure 4:**
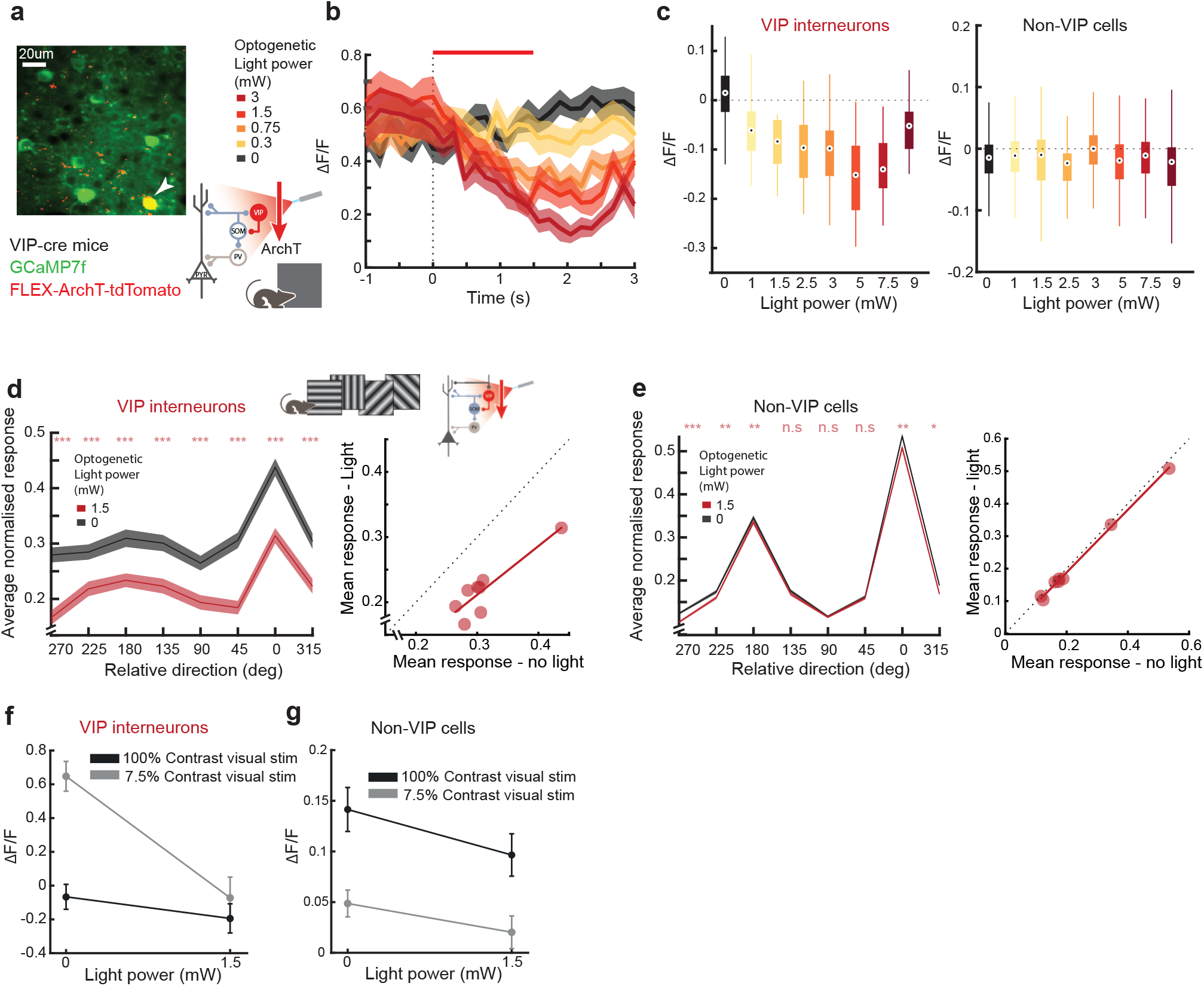
VIP inhibition moderately suppresses responses to visual stimuli during passive presentation. **a**) Example region of an *in vivo* imaged plane showing all neurons expressing GCaMP7f and a VIP interneuron (arrowhead) additionally expressing ArchT-tdTomato. **b**) Mean responses of an example VIP interneuron to different light powers. Responses are aligned to optogenetic light onset (dashed line). Red bar indicates optogenetic stimulation duration (1.5s), shading indicates SEM. **c**) Box plots of optogenetically inhibited activity (mean 0-1.5s, baseline subtracted) across different light powers for VIP interneurons (left, n = 92 cells) and non-VIP cells (right, n = 1648 cells, 5 mice). **d**) Left, stimulus evoked normalised activity in response to different oriented gratings, averaged across all VIP interneurons (n = 122 cells, 6 mice) aligned to their preferred direction. *, p<0.05, **, p<0.01, ***, p<0.001 Wilcoxon signed-rank test for photoinhibition compared to non-photoinhibition conditions at each direction, corrected for multiple comparisons. Right, the same data shown as average activity with and without optogenetic light. Linear regression, slope = 0.748, intercept = -0.012. **e**) Same as d, for all orientation selective non-VIP neurons, n = 953 cells, slope = 0.965, intercept = -0.005. High power, slope = 1.634, intercept = 0.032, 8 mice. **f**) Visual stimulus evoked VIP interneuron activity (mean 0-1.5s, baseline subtracted) in response to a drifting vertical grating at low and high contrast, with and without VIP photoinhibition, n = 37 cells, 5 mice. **g**) Same as h, for non-VIP cells, n = 528 cells, 5 mice.

We photoinhibited VIP interneurons while passively presenting drifting visual grating stimuli to the mice (Fig. 4d, left). VIP interneurons decreased activity at each grating direction with photoinhibition (Fig. 4d, right, slope = 0.748, intercept = -0.012, n = 122 cells). As a result, stimulus responses of orientation selective non-VIP cells were moderately but significantly inhibited at specific directions, including at the peaks of the tuning curve (Fig. 4e, slope = 0.965, intercept = -0.005, n = 953 cells). To confirm that we had inhibited VIP interneurons to minimal activity levels, we tested another group of mice in which we presented low contrast grating stimuli which activate VIP interneurons more strongly than high contrast stimuli ^50^. When we combined stimulus presentation with VIP photoinhibition, VIP interneuron activity was reduced to the same level whether they were highly active or less active (Fig. 4f). VIP inhibition in both cases was accompanied by reductions in non-VIP cell activity (Fig. 4g). This confirmed that our photoinhibition was appropriate for reducing VIP activity to near the minimum physiological levels. Thus, VIP inhibition during passive viewing of visual stimuli led to a modest inhibition of non-VIP cell activity. However, to establish whether VIP interneurons are required for attentional modulation of stimulus selectivity we next studied the effects of inhibition of VIP activity during the attention-switching task.

### VIP inactivation during attention-switching

To conclusively rule out the role of VIP interneurons in attentional modulation of stimulus selectivity, we photoinhibited VIP interneurons in randomly interleaved trials while mice performed the attention-switching task, with the light on period from 100ms before the grating stimulus onset until the end of the stimulus (Fig. 5a). As with photoactivation, photoinhibition had no effect on the mouse behaviour (Supplementary Fig. 3c, d). VIP interneurons were inhibited during the behaviour, both when the visual stimuli were attended and ignored (Fig. 5b, top). However, we observed no effect of VIP inhibition on stimulus responses of non-VIP neurons (Fig. 5b, bottom). To confirm that there was no deficit in the degree of our photoinhibition, we applied an average of 7.5 mW light power (range 5.7 to 9 mW), higher than the 1.5 mW used so far. We found no further reduction in VIP interneuron activity at this higher light power (Supplementary Fig. 4a). Furthermore, we found that the higher light power led to a significant increase in measured neural activity in the tdTomato control mice (Supplementary Fig. 4b). This revealed the potential for artefactual results with the higher light power, possibly induced through direct retinal activation and subtle changes in behaviour linked to light onset (Supplementary Fig. 4c, d). These results illustrate the importance of light-only controls in animals performing the full behavioural task and confirmed that our photoinhibition of VIP interneurons at 1.5 mW was close to complete.

**Figure 5:**
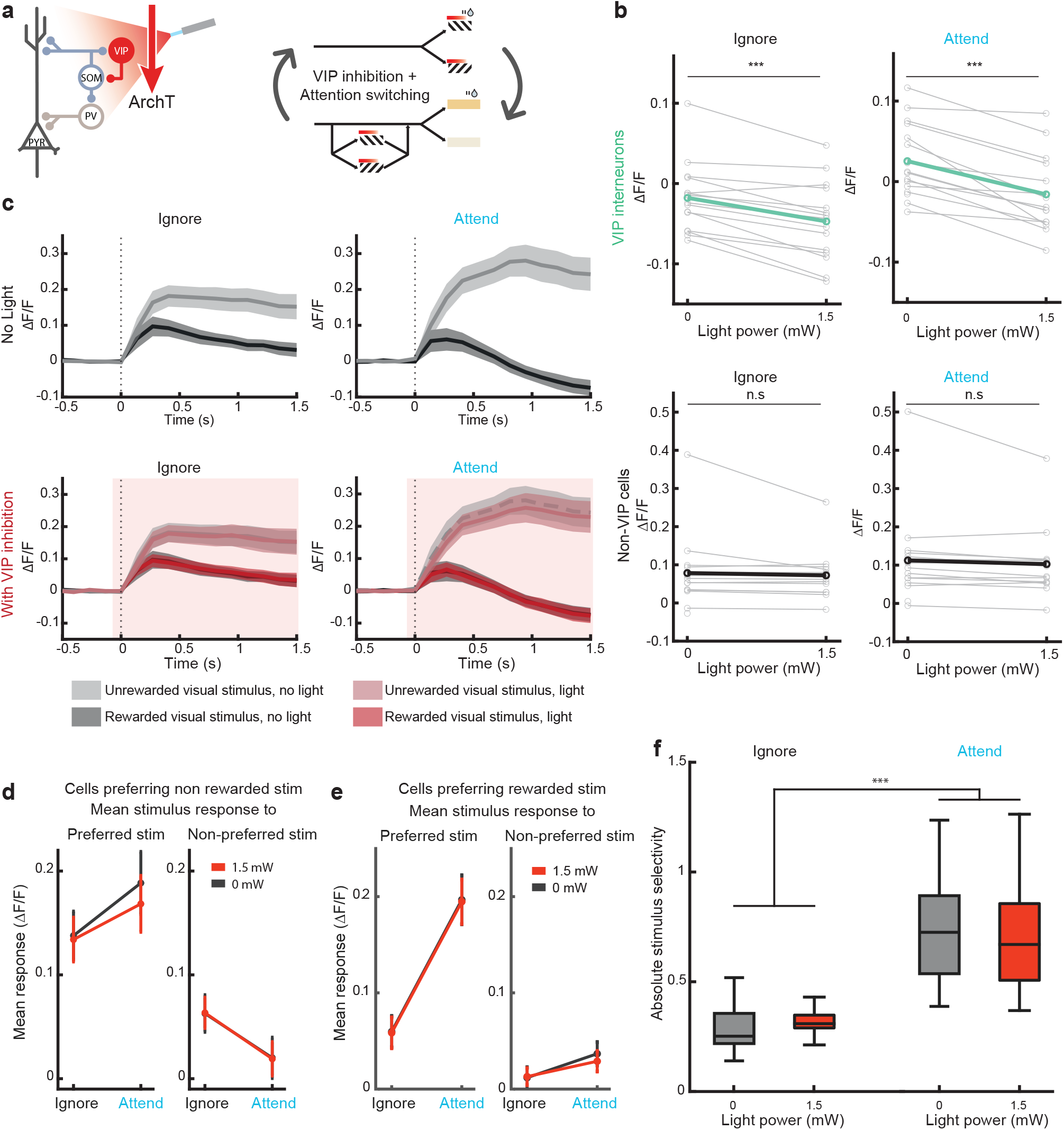
No effect of VIP inhibition on attentional modulation. **a**) Schematic showing VIP photoinhibition during the attention switching task. Light onset (red bars) was from -0.1s to 1.5s relative to visual stimulus onset. Light was ramped off over 0.2s. **b**) Mean visual stimulus evoked activity (baseline subtracted) with increasing VIP photoinhibition. Top, all VIP interneurons, bottom, all non-VIP cells. Left, responses when ignoring the visual stimuli, right, responses when attending the visual stimuli. Wilcoxon signed-rank test between photoactivation and non-photoactivation conditions, ***, p< 0.001, n = 16 sessions, 5 mice. Gray lines indicate individual session averages, coloured lines indicate overall average. **c**) Top, mean visual stimulus evoked activity for all non-VIP cells with preference for the unrewarded stimulus that significantly increased their selectivity with attention (mean of n = 16 sessions, shading indicates SEM). Bottom, same sessions, responses with additional VIP photoinhibition (red). Responses from top are superimposed for comparison (grey dashed lines, light red shading indicates light onset). **d**) Black, mean visual stimulus evoked activity (averaged 0-1s, baseline subtracted) of all non-VIP cells with preference for the unrewarded stimulus that significantly increased their selectivity with attention, n = 16 sessions. Red, same responses with additional VIP photoinhibition. Error bars indicate SEM. **e**) Same as d for cells with preference for the rewarded stimulus. **f**) Absolute stimulus selectivity without and with VIP photoinhibition for all non-VIP cells which significantly increased their selectivity with attention (n = 16 sessions). Stimulus selectivity measured when ignoring the visual stimuli (left) and attending the same stimuli (right). There was a significant effect of attention on selectivity, but not of VIP activation or an interaction between the two (2-way ANOVA, attention p = 1.13×10^−16^, other ps > 0.05).

To study the interaction of VIP inhibition with attention, we selected all non-VIP neurons which showed a significant increase in stimulus selectivity with attention. The response of this population to the two grating stimuli in the ignore and attend conditions, with and without VIP photoinhibition showed no effect of VIP inhibition (Fig. 5c, average of all cells with preference for unrewarded stimulus). Increasing light power had no effect on the average (0 to 1s) activity of this population in either ignore or attend conditions, for either the preferred or non-preferred stimulus (Fig. 5d). We performed a two-way ANOVA for the rewarded stimulus responses and found a significant main effect for attention (p = 0.001) but not for VIP inhibition (p = 0.80), and no significant interaction effect (p = 0.97). Similar results of no interaction between attention and VIP inhibition were obtained for the unrewarded stimulus and from cells with a preference for the rewarded stimulus (Fig. 5e).

Having found no effect of VIP inhibition on non-VIP cell stimulus evoked activity, we asked whether there was any change in stimulus selectivity with VIP inhibition in either the ignore or attend condition (Fig. 5f). We performed a two-way ANOVA on the data in Fig 5f (all non-VIP cells which significantly increased their selectivity with attention) and found a significant main effect on selectivity from attention (p = 1.13×10^−16^), but not from VIP activation (p = 0.88) and no interaction effect (p = 0.63). The same result was obtained when taking all non-VIP cells (a significant main effect on selectivity from attention, p = 1.72×10^−10^, but not from VIP activation, p = 0.89 and no interaction effect, p = 0.73). Thus, VIP interneurons are not involved in producing the gain in stimulus selectivity during attention.

Overall, although VIP interneurons are capable of strongly modulating activity of the non-VIP neural population, they are not the route through which attention induces selectivity changes in V1.

### Attention and VIP modulations are orthogonal

Although VIP disinhibition is not involved in attention modulation, can attention-driven and VIP-driven modulations co-occur in the same neurons and yet have no adverse impact on each other? To address this question, we asked if these two modulations were orthogonal to each other. We performed dimensionality reduction using linear discriminant analysis (LDA) and found two axes best separating the visual stimulus-evoked neural activity of the non-VIP population: first between attend and ignore conditions, and second between photoactivation and no photoactivation conditions (VIP activation, VIP inhibition and control) within the ignore condition (Fig. 6a). We found the cosine similarity of these two axes as a measure of the alignment of the attentional and optogenetic manipulations.

**Figure 6:**
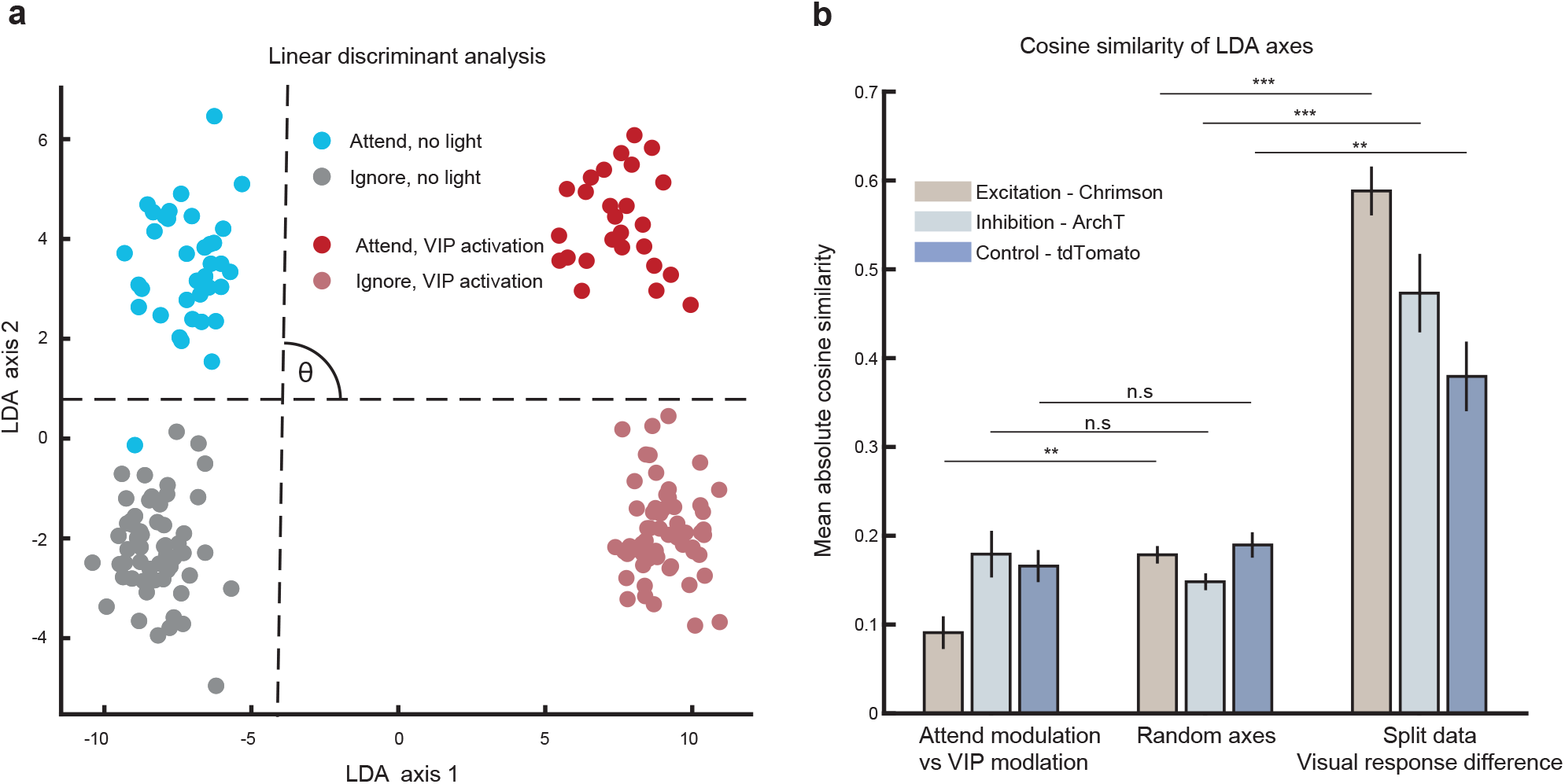
Attention and VIP modulations are orthogonal **a**) Mean stimulus evoked activity from individual trials projected onto the first two axes obtained from dimensionality reduction using linear discriminant analysis (LDA). Example session from a mouse expressing Chrimson. Dashed lines indicate the directions along which attention and VIP photoactivation most strongly modulated activity. **b**) Mean cosine similarity for pairs of axes in neural activity space (Chrimson n = 17 sessions, ArchT n = 16 sessions, tdTomato control n = 17 sessions). From left to right, absolute cosine similarity for: LDA axis separating VIP photoactivation vs no photoactivation trials and LDA axis separating attend vs ignore trials; pairs of random axes extracted based on the covariance of the original neural data (10,000 samples); two axes separating the rewarded vs unrewarded visual stimuli where each axis was found using half of the data (mean of 50 shuffled repeats). Significance tests were against the corresponding random axes median, ** p<0.01, *** p<0.001. From left to right p values for all tests: 0.009, 0.379, 0.653, 6.43e-04, 2.93e-04, 0.002.

For comparison, we created a null distribution of cosine similarity between randomly chosen pairs of axes from the dataset based on the covariance of the original data (10,000 samples, same method as^51^). We compared the cosine similarity of the LDA axes of modulation by attention and modulation by VIP manipulation for each session to its random mean. In the VIP activation experiment, the angle between attention and VIP modulations had a significantly lower cosine similarity value than the random vectors (Fig. 6b, Wilcoxon signed-rank test, p = 0.009, n = 17 sessions), demonstrating that they were significantly less aligned than random axes drawn from the same space occupied by the data. This demonstrates that attention and VIP disinhibition produce modulations that are significantly closer to orthogonality than expected by chance^51^.

The VIP inhibition and control experiments showed no difference in cosine similarity from random vectors (Fig. 6b, Wilcoxon signed-rank test, p = 0.4, n = 16 sessions, p = 0.7, n = 17 sessions respectively), consistent with the absence of light-driven effects in these two groups. As a positive control, we confirmed that in all three groups, the vectors separating the two visual stimuli responses from two halves of the data had significantly higher cosine similarity than random vectors (Fig. 6b, Wilcoxon signed-rank test Chrimson p = 2.93×10^−04^, ArchT p = 6.43×10^−04^, tdTomato p = 0.002. n = 16, n = 17, and n = 17 sessions respectively). These results demonstrate that not only do attention and VIP modulation not interact, but they are also orthogonal to each other.

### Distinct mechanisms of changes during attention and VIP modulation

Having established that VIP-induced and attention-induced modulations do not interact and are in fact orthogonal, we wished to understand the circuit mechanisms underlying these two modulations and to establish how similar or distinct they were. Although cortical VIP interneurons are known to strongly inhibit SOM interneurons and thus disinhibit pyramidal neurons, these cells are embedded in highly recurrently connected networks with strong connectivity across cell classes^17,19^. It is thus often not clear how the perturbation of one cell class influences the rest of the network^52–54^. In fact, the effect of VIP activation leading to SOM inhibition and thus PYR disinhibition has never been demonstrated in simultaneously measured VIP, SOM and PYR cells *in vivo*.

To establish the cell class-specific consequences of our optogenetic VIP activation *in vivo*, we reidentified the same neurons in co-registered, immunohistochemically stained brain sections^5,55^ in 4 out of 8 of the same animals expressing Chrimson in VIP interneurons, and detected simultaneously imaged PV, SOM and VIP positive interneurons (Fig. 7a). The remaining cells were classified as putative PYR cells. We could thus measure the effect of VIP photoactivation on the activity of multiple simultaneously measured cell classes. VIP photoactivation during passive grey screen viewing led to increasing disinhibition of PYR and PV cells with increasing light power (Supplementary Fig. 5a-e). VIP photoactivation during passive presentation of oriented drifting grating stimuli showed, as expected, a strong enhancement of VIP and PYR cell responses. Crucially, at the same time we saw a suppression of SOM cell activity on average, and an enhancement of PV cell activity (Fig. 7b, c, d). This is consistent with VIP photoactivation leading to disinhibition of PYR and PV cells via direct inhibition of SOM interneurons.

**Figure 7:**
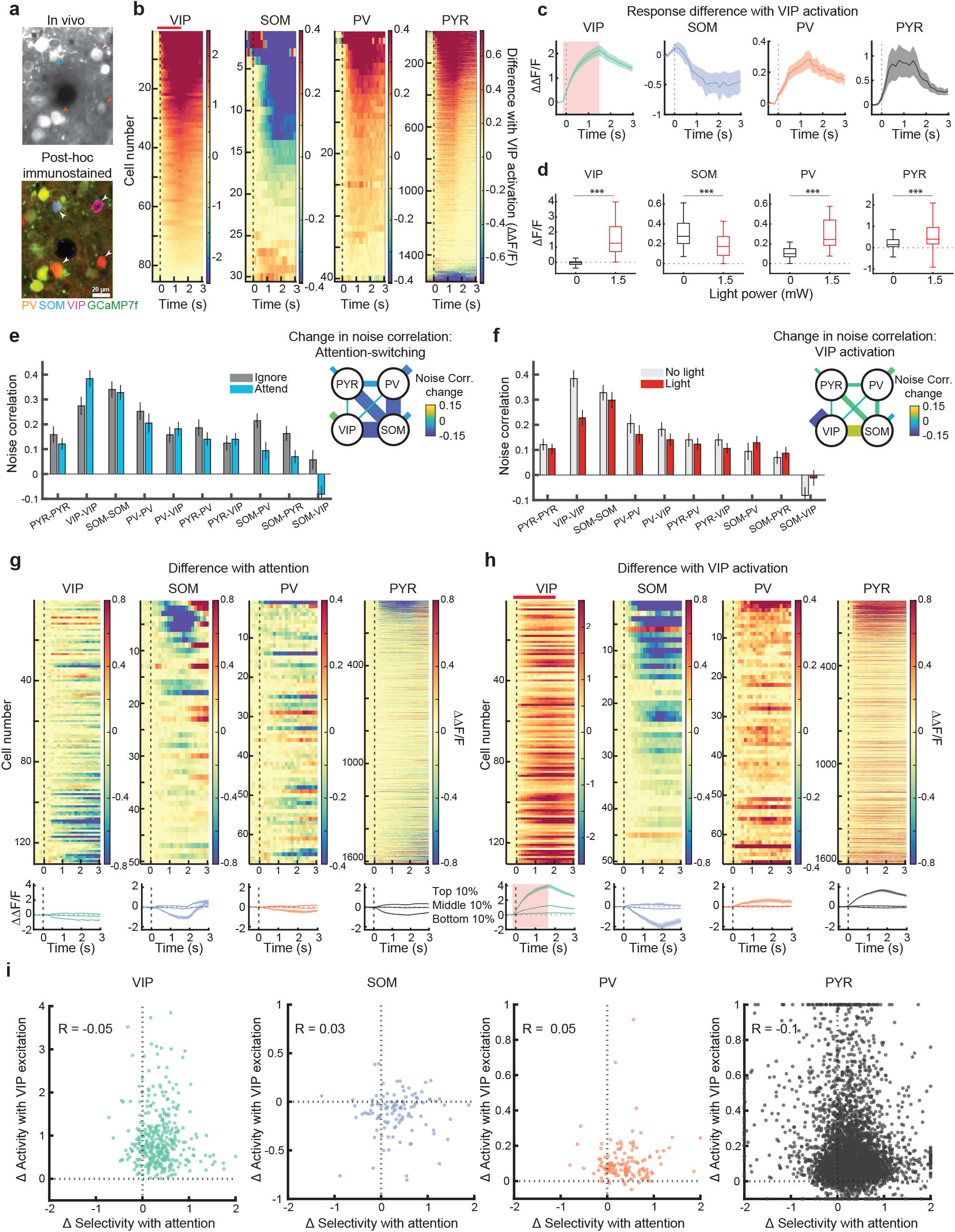
Simultaneous VIP, SOM, PV and pyramidal cell activity reveals distinct mechanisms of modulation with attention and VIP photoactivation **a)** Example region of an *in vivo* image plane with GCaMP7f-expressing neurons (top) and the same region after post hoc immunostaining for PV, SOM, and VIP (orange, blue, and magenta, respectively) following image registration (bottom). Identified interneurons are indicated by arrowheads. **b**) Difference (VIP photoactivation condition minus no photoactivation condition) of average visual stimulus-evoked response for each cell in all 4 cell classes (average of all orientations of visual stimuli during passive presentation) aligned to visual stimulus onset (dashed line). Optogenetic light onset here and below is -0.1s to 1.5s from visual stimulus onset (red shading). Cells are sorted by their average response difference 0–1 s from stimulus onset. Left to right: VIP n = 85 cells, SOM n = 30 cells, PV n = 40 cells, PYR n = 1567 cells. **c**) Mean of each column of b, showing average change in activity with VIP activation, shading indicates SEM. **d**) Box plots of visual stimulus evoked activity with and without VIP photoactivation, averaged 0 to 1s from visual stimulus onset, for each cell class (Wilcoxon signed-rank tests for differences in activity: PYR, n = 1567 cells, p = 4.52×10^−187^; SOM, n = 30 cells, p = 5.71×10^−04^; VIP, n = 85 cells, p = 1.17×10^−15^; PV, n = 40 cells, p = 3.57×10^−08^). **e**) Mean noise correlations between cell pairs belonging to the same or different cell classes, in the ignore and attend conditions, without photoactivation. Error bars represent SEM here and below (n = 11 sessions, 4 mice). Inset: Changes in noise correlations due to attention as indicated by line thickness and colour code. Shorter line segments indicate change in noise correlations between cells of the same type. **f**) Same as e, for noise correlations with and without VIP photoactivation in the attend condition. **g**) Top, difference in mean visual stimulus evoked response with attention (baseline subtracted, unrewarded grating), for each cell type, aligned to visual stimulus onset (dashed line). Cells are sorted by their average activity in the ignore condition (see also Supplementary Fig. 6). Bottom, average responses of cells from the top, middle and bottom 10th percentiles of the difference in responses (averaged 0-1s) with attention shown above. Shaded area indicates SEM. **h**) Same as g but for differences in mean visual stimulus evoked response with photoactivation of VIP interneurons (red bar and shading) compared to no photoactivation in the ignore condition. Top, cells are sorted the same as in g, by their average activity in the ignore condition. **i**) Relationship between ΔSelectivity with attention (positive values indicate increased stimulus selectivity with attention) and change in stimulus evoked activity with VIP photoactivation (mean 0-1s, baseline subtracted), for VIP (N = 130 cells), SOM (N = 50 cells), PV (N = 67 cells) and PYR cells (N = 1616 cells, 4 mice). Cells with values greater than the axes limits were pegged to the axes. Significant negative correlations were present only in VIP and PYR cells.

The direction tuning curves of each cell class measured during passive viewing were also modulated following a similar pattern, with PYR (high power, slope = 1.452, intercept = 0.042, n = 472 cells), PV (slope = 1.304, intercept = 0.125, n = 40 cells) and VIP (slope = 1.489, intercept = 0.567, n = 85 cells) cell activity largely enhanced and SOM (slope = 0.690, intercept = 0.019, n = 30 cells) cell activity largely inhibited at all stimulus directions during VIP photoactivation (Supplementary Fig. 5f-j). These results demonstrated that VIP interneuron activation drives network-wide changes in all major cell classes and confirmed that activating VIP interneurons within physiological levels leads to disinhibition of PYR cells through SOM inhibition. The enhanced activity of PV cells with VIP activation demonstrated that the direct inhibition of PV cells by VIP cells was reversed by other network effects, which we explored in a circuit model below.

If attention and VIP modulations are indeed orthogonal, we would expect the underlying mechanisms of changes induced by either of them to be distinct. We tested this using different measures. First, as shown above, attention led to both increases and decreases in average stimulus evoked activity (Fig. 1i), but VIP activation only induced increases in stimulus evoked activity (Fig. 3b, d). Second, we measured noise correlations between the 4 cell classes we had recorded from simultaneously. Noise correlations were measured as the stimulus-independent trial-to-trial co-variability of responses, and thus provided an estimate of mutual connectivity and shared inputs between and within cell classes. We found that attention significantly increased noise correlation between VIP cell pairs and significantly decreased the correlation between PYR and SOM, SOM and VIP and PV and SOM cell pairs (Fig. 7e). In contrast, VIP activation significantly decreased noise correlation between PYR, VIP and PV cell pairs as well as between PYR and VIP, PV and VIP and SOM and VIP cell pairs (Fig. 7f). P-values (uncorrected, unrewarded visual stimulus) for changes with attention are PYR-PYR: p = 0.320, VIP-VIP: p = 0.005, SOM-SOM: p = 0.966, PV-PV: p = 0.123, PV-VIP: p = 0.278, PYR-PV: p = 0.054, PYR-VIP: p = 0.123, PV-SOM: p = 0.014, PYR-SOM: p = 0.019, SOM-VIP: p = 0.005; for changes with VIP photoactivation, PYR-PYR: p = 0.010, VIP-VIP: p = 0.001, SOM-SOM: p = 0.240, PV-PV: p = 0.019, PV-VIP: p = 0.010, PYR-PV: p = 0.067, PYR-VIP: p = 0.032, PV-SOM: p = 0.083, PYR-SOM: p = 0.278, SOM-VIP: p = 0.003.

Third, we studied how attention and VIP photoactivation modified stimulus evoked activity across the populations of these 4 cell classes. We found a heterogeneity in the responses to visual stimuli in all cell classes (Supplementary Fig. 6). By measuring the difference in stimulus evoked responses between attend and ignore conditions for each neuron, we found that in the absence of photoactivation, attention led to heterogeneous changes, marked by a prominent reduction in evoked activity in all four cell classes (Fig. 7g). In contrast, VIP photoactivation led to predominantly enhanced activity in VIP, PV and PYR neurons, and largely suppressed activity in SOM neurons (Fig. 7h).

Fourth, we directly compared the degree of selectivity modulation by attention and activity modulation by VIP activation on neurons from each cell class (Fig. 7i). We found no strong correlation in any cell class, ruling out the possibility of a specific subset of cells driving both attentional and VIP-driven changes (Pearson’s correlation coefficients -0.05, 0.03, 0.05, -0.1 in VIP, SOM, PV and PYR cells respectively). In fact, we found a small but significant negative correlation in PYR (p = 3.4×10^−6^) and VIP interneurons (p = 0.015, all other ps > 0.05), suggesting a moderate segregation of attention and VIP modulations across these cell populations. Overall, these results demonstrate that attention changes V1 stimulus processing by heterogeneous changes in activity and correlations across different cell classes, whereas VIP activation drives relatively homogenous enhancement of PYR and PV cell activity along with SOM inhibition. Thus, attention and VIP disinhibition act on the same cortical circuit through distinct mechanisms.

### A circuit model captures the effects of both attention and VIP perturbations

Our results raised an important question regarding cortical circuit architecture: How can attention and VIP modulation each profoundly affect the same cell populations and yet not interact? Cortical circuits have structured connections between cell classes and receive targeted bottom-up and top-down inputs. What circuit architecture can account for increased stimulus selectivity through top-down attentional inputs, modulation of PYR, PV and SOM cells through VIP activation, and no interaction between these two effects? To address this question, we developed a theoretical model of the excitatory and inhibitory circuit enabling us to determine what patterns of connectivity and external inputs could explain the results of both attentional and VIP-driven modulations of the same circuit.

In this model, we represented the four cell types PYR, PV, SOM, and VIP by their population activity. The activity of each population was determined by baseline activity, bottom-up stimulus-related input, top-down attentional input, and connection strengths with other cell populations (Fig. 8a). We used experimentally derived connectivity values from ^17^, and used simulation-based inference (SBI)^56^ to determine connectivity values from PYR to all interneurons, SOM to PV cells, bottom-up, top-down and baseline inputs which replicated a number of experimental findings, importantly that of non-interacting attention and VIP modulations (see methods).

**Figure 8:**
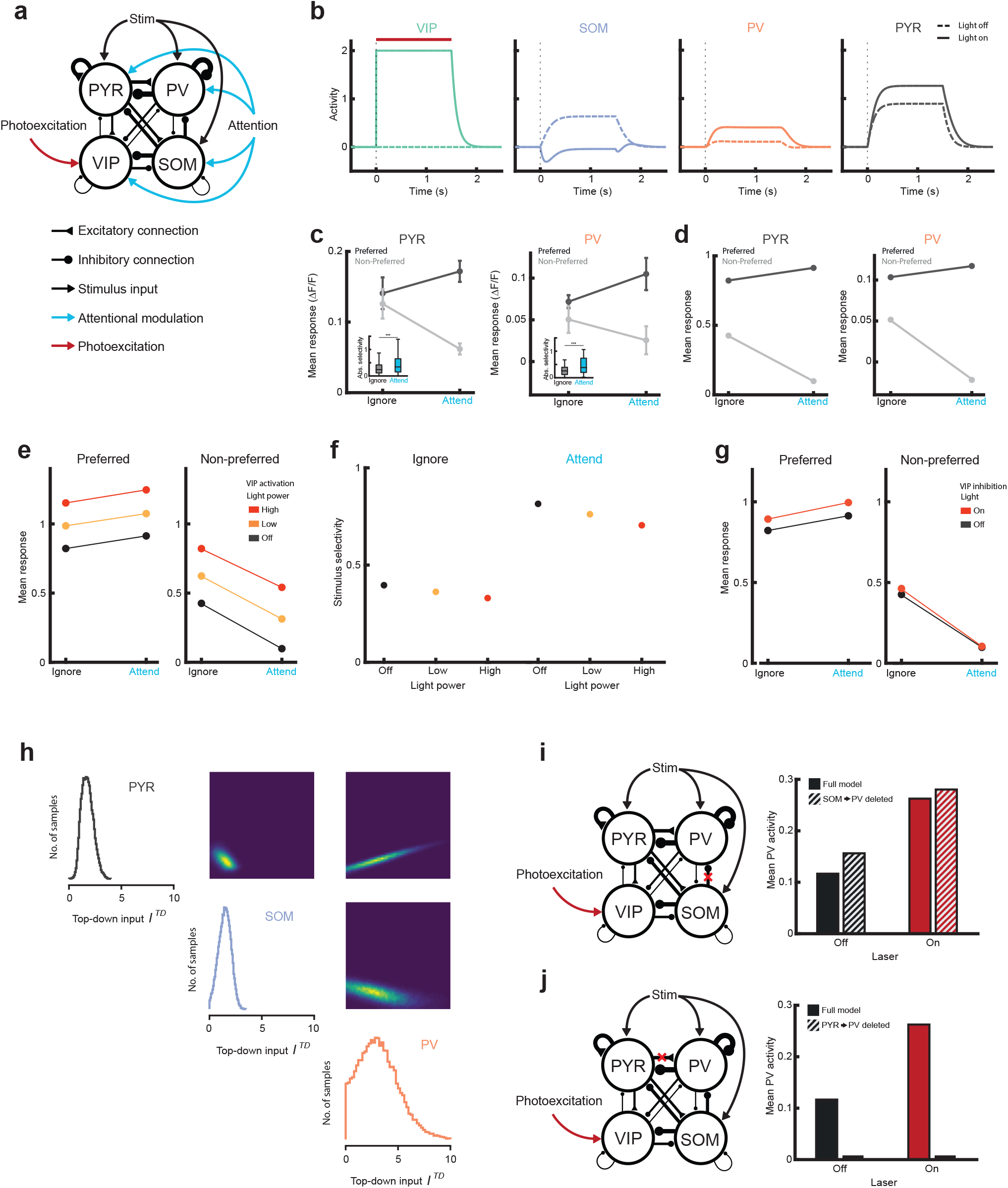
Circuit modelling reveals a network architecture for independent attentional and VIP modulations **a**) Schematic of the model architecture, indicating connectivity between different cell classes, bottom up (stim) and top-down inputs (attention), and VIP photoactivation. **b**) Simulated responses of the 4 cell types to a visual stimulus with and without VIP photoactivation. **c**) Experimental results, average responses to the preferred and non-preferred visual stimuli in the ignore and attend conditions for positively selective cells that significantly increased their stimulus selectivity with attention, for PYR (left, n = 289 cells,) and PV cells (right, n = 15 cells), error bars indicate SEM. Inset: absolute selectivity of the full population of PYR (n = 1616) and PV cells (n = 67), Wilcoxon signed rank test, PYR p = 7.02×10^−21^, PV p = 8.53×10^−5^. **d**) Model output, same as c. **e**) Mean response of the model PYR population to visual stimuli with and without attention, and with no, low or high VIP photoactivation, for the preferred (left) and non-preferred stimuli (right). Compare to Fig. 3d. **f**) Stimulus selectivity of the model PYR population with and without attention, and with no, low or high VIP photoactivation. Compare to Fig. 3f. **g**) Same as e, for VIP photoinhibition. Compare to Fig. 5d, e. **h**) Visualization of the posterior distribution over the three parameters for the top-down modulation of PYR, SOM, and PV which were consistent with the experimental observations, and obtained by sampling 50,000 parameter sets from the estimated posterior distribution (obtained using the Sequential Neural Posterior Estimation algorithm from the simulation-based inference package). Shown are the univariate marginals (histograms) and pairwise marginals (2-dimensional histograms). The y-axis of each 2-dimensional histogram corresponds to the values of the parameter on the left of it, and the x-axis to the values of the parameter below. The colour indicates the number of samples in each bin. **i**) Model in which the SOM to PV connection was set to zero (left). The model responses of the PV population were compared, between the full model and model with the SOM to PV connection deleted, with and without VIP photoactivation (right). No strong changes were observed. **j**) Same as i, with PYR to PV connections set to zero. PV responses were strongly disrupted and did not change with VIP photoactivation.

We confirmed that the model reproduced the suppression of SOM and enhancement of PYR and PV activity during VIP activation (Fig. 8b). We next compared activity with and without attentional top-down inputs, focusing on PYR and PV cells since these cells showed a robust increase in stimulus selectivity, uninfluenced by overt movements (PYR average absolute selectivity, ignore 0.43 ± 0.77 [mean ± SD], attend 0.49 ± 0.52, Wilcoxon signed rank test, p = 7.02×10^−21^, N = 1616. PV, ignore 0.29 ± 0.23, attend 0.48 ± 0.34, p = 8.53×10^−5^, N = 67. SOM, ignore 0.35 ± 0.33, attend 0.45 ± 0.34, p = 0.08, N = 50. VIP, ignore 0.23 ± 0.17, attend 0.41 ± 0.30, p = 7.89×10^−09^, N = 130. Fig. 8c. See also ^5^). The model reproduced the increase in PYR and PV stimulus selectivity through a combination of increased and decreased responses to the preferred and non-preferred stimuli respectively (Fig. 8d). Crucially, the model captured the non-interacting nature of attention and VIP modulations. VIP activation enhanced PYR activity almost equally in the ignore and attend conditions (Fig. 8e), closely matching the experimental findings (Fig. 3d). This led to small changes in stimulus selectivity with VIP activation, but large selectivity changes with attention (Fig. 8f, compare to Fig. 3e). The model responses to VIP inhibition (Fig. 8g) also closely followed experimental results (Fig 5d-e), even though this observation was not used to constrain the model. The model thus revealed a precise network architecture capable of sustaining strong yet non-interacting modulations by attention and VIP activation.

Our model allowed us to ask which cell class might receive the strongest top-down inputs to account for the data. We explored the parameter distribution which satisfied the data and found that PV cells received the strongest top-down inputs (Fig. 8h). Furthermore, top-down inputs arriving on PYR cells were tightly correlated with those arriving on PV cells (Fig. 8h), suggesting that the relative strength of top-down inputs on PYR and PV cells is highly regulated.

Finally, we explored the origin of the experimental finding that VIP activation led to enhanced activity not only in PYR cells but prominently in PV cells as well (Fig 7b-d). The enhancement of PV cell activity with VIP activation may be a result of disinhibition through SOM cells, since SOM cells inhibit PV cells^17^, or a consequence of elevated PYR activity leading to increased PV cell activity through PYR to PV connections ^57^, or a combination of the two effects. Our circuit model allowed us to test which of these scenarios was more likely. In our model, we first set the connectivity weight from SOM to PV cells to zero and compared the response of PV cells with and without VIP activation. We found a minimal effect of deleting the SOM to PV connection (Fig 8i). In contrast, when we deleted the connection from PYR to PV cells we found strongly reduced PV responses which were not affected by VIP activation (Fig 8j). These results suggest that VIP activation first suppresses SOM cells, leading to enhanced PYR activity due to disinhibition, which subsequently increases PV activity through PYR to PV connections.

## Discussion

In this study we demonstrate that attention leads to modulation in stimulus response properties of V1 neurons resulting in increased stimulus selectivity. We further demonstrate that VIP-SOM disinhibition leads to strong modulations in V1 stimulus response properties. Critically, however, these two modulations do not interact, and are in fact orthogonal to each other at the population level. Inhibition of VIP interneurons leaves attentional modulation unperturbed, and VIP interneurons themselves are not modulated by attention, confirming the lack of involvement of VIP interneurons in generating attentional modulation. Consistent with this conclusion, attention and VIP modulations are accompanied by distinct patterns of changes of activity and interactions between excitatory and different inhibitory cell classes. Circuit modelling revealed a precise network architecture compatible with these experimental findings.

### Circuit basis of attentional modulation

Recent studies of visual attention in mice have allowed the use of genetic, molecular, viral and optical techniques to obtain detailed circuit-level insights into this key cognitive phenomenon^7,47,58–61^. This has provided insights into the role of specific molecularly defined cell classes, and their targeting by long range, top-down projections^38,62,63^. However, it is challenging to test hypotheses about the circuit basis of cognitive phenomena since this requires the measurement and manipulation of neural circuit components as animals perform a cognitive task. In this study we have aimed to advance this approach, by simultaneously measuring the activity of four cell classes while manipulating the activity of one of these cell classes, as animals switched between distinct attentional states. Using this approach, we were also able to calibrate our circuit manipulations to ensure they spanned a physiologically relevant range. As a result, we established that VIP cells are not the route through which top-down attentional signals influence activity in V1.

An alternative explanation may be that a specific pattern of individual VIP cell activation needs to be engaged by attentional signals, and our non-specific full-field photoactivation approach is insufficient to rule out the role of VIP interneurons in attention modulation. However, this argument is refuted by our findings that VIP interneuron inhibition resulted in no changes to attentional modulations, and that VIP interneuron responses themselves were not modulated by attention. What alternative local circuits might then be involved in implementing the observed attentional modulations? One possibility is that attention induces selective suppression through PV interneurons, which themselves show strong attentional modulations^5,64^, and may be capable of enhancing stimulus selectivity and discrimination behaviour^65^. This may occur in conjunction with direct excitatory top-down projections onto pyramidal neurons, leading to enhanced activity during attention^63,66^.

### Role of VIP disinhibition

What is VIP-SOM disinhibition in V1 involved in if not attention? The effect of local visual context (stimulus surround) may act through a VIP-SOM disinhibitory circuit^67^. This circuit has also been strongly implicated in running-induced modulations in V1^29^, which has parallels with attentional gain modulation^1,47^ but appears mechanistically distinct^8^. However, more than the immediate effect on network activity, VIP-SOM driven disinhibition may be crucial in gating plasticity onto pyramidal cells^24^, allowing for associative^68^ or motor learning^69^.

Given the richly interconnected network of excitatory and inhibitory cortical cell classes^17,19^, it was not surprising that activation of VIP interneurons led to robust changes in stimulus responses in all other simultaneously measured cell classes. These network effects were dominated by inhibited SOM activity and elevated activity in PYR and PV cells. Although this pattern was observed on average, the effects of photoactivation and attention revealed heterogeneity within cell populations, which is not surprising given that each molecularly defined class studied here comprises sub-types with distinct gene expression patterns, morphologies, network connectivity and intrinsic properties^18^.

The motif of VIP activation leading to PYR disinhibition through SOM inhibition, appeared in various stimulus evoked conditions including orientation mapping (with no behavioural task) and during the full behavioural task. However, this interpretation runs into challenges when considering the effect of VIP activation on other cell classes without visual stimuli (Supplementary Fig. 5). In this condition, VIP activation led to inhibition of only a small fraction of SOM cells, yet led to robust elevation of PYR and PV activity. This scenario has parallels with the findings by ^52^ comparing running evoked disinhibition in the dark and with visual stimulation. While those results could be explained by a theoretical model incorporating a nonlinearity in SOM response functions^53^, our dataset may not fit such a model due to the matched luminance levels across all conditions.

In general, caution is required in interpreting any results of circuit manipulations in densely interconnected and active networks^54^. In our study, however, several findings converge to the same conclusion: the absence of modulation of VIP cells by attention, the absence of interactions between attention and VIP activation effects, the absence of any impact with VIP inhibition, and the orthogonality of the two manipulations at the population level, all support the conclusion that the two modulations are independent.

### Orthogonal modulations

Orthogonal representations are important in segregating movement-related signals from stimulus evoked activity in visual cortex^70^. In the primate motor cortex during reaching movements, preparatory activity before the reach is actively maintained orthogonal to the activity during the reach, possibly supporting non-interfering computations^51^. In the mouse somatosensory cortex, approximately orthogonal representations of whisker contacts may allow flexible use of the same stimulus representations in distinct tasks^71^. Given that we found that attentional signals are orthogonal to VIP modulations, what distinct computations might be multiplexed using this property? Since VIP interneurons in V1 do receive top-down inputs from prefrontal cortex, other prefrontal signals may take advantage of this orthogonality, to additionally engage the same V1 populations in other cognitive processes along with attention, such as working memory^72^, or prediction error computation^73^.

Overall, our results suggest the need to revise our understanding of the role of VIP-SOM driven disinhibition in attentional modulation of stimulus selectivity. At the same time, we demonstrate the remarkable capacity of cortical circuits to combine multiple orthogonal and mechanistically distinct computations in the same neural populations.

## Author contributions

DM-J and AGK designed the experiments. DM-J performed the experiments and analysed the data. MF-O provided technical assistance and performed the immunolabelling and post-hoc cell matching. KW developed and analysed the circuit model, with inputs from CC. AGK and DM-J wrote the paper with inputs from KW and CC. All authors contributed to discussions and commented on the manuscript.

## Acknowledgments

We thank Florencia Iacaruso, Takahiro Kanamori, Dimitar Kostadinov and Petr Znamenskiy for discussions and comments on the manuscript, and Matthew Grubb, Samuel Cooke, Juan Burrone, Attila Losonczy and members of the Khan laboratory for discussions of the results in this manuscript. This work was supported by the Wellcome Trust (AGK, 206222/Z/17/Z), the BBSRC (AGK BB/S015809/1), and start-up funds from the CDN, King’s College London (AGK).

## Materials and methods

All experimental procedures were carried out in accordance with institutional animal welfare guidelines and licensed by the UK Home Office.

### Animals and surgical procedures

A total of 20 VIP-Cre mice (MGI:4431361) were used in this study (5 male, 15 female). Mice aged between postnatal days 78 to 84 were anaesthetised using isoflurane, at 4% concentration for induction and at 1-1.5% for maintenance. At the start of the surgery additional drugs were given to provide analgesia (Metacam 5mg/kg), anti-inflammatory effects (dexamethasone 3.8mg/kg), and to reduce mucus secretions (Atropine 0.08mg/kg). Eye-cream (Maxitrol) was applied to the eyes to prevent drying and body temperature was maintained at 37°C using a heating mat and rectal temperature probe (Harvard Apparatus).

Mice were implanted with a chronic imaging window (3-5 mm diameter) above the right V1 (2.7mm medial, 0.6mm anterior to lambda) following viral injections using a pressure micro-injection system (Picospritzer III, Parker) of AAV1-hSyn-GCaMP7f mixed with either AAV5-hSyn-FLEX-ChrimsonR-tdTomato (8 mice), AAV5-hSyn-FLEX-ArchT-tdTomato (5 mice) or AAV1-FLEX-tdTomato (3 mice). Mice used in the low contrast stimulus experiments were injected with a mixture of AAV8-hEF1a-jGCaMP7f and AAV5-hSyn-FLEX-ArchT-tdTomato (4 mice). The craniotomy was then sealed with a glass coverslip and cyano-acrylic glue (Loctite). A custom machined aluminium head-plate was cemented onto the skull using dental cement (C&B Superbond).

Before the removal of anaesthesia mice were injected with antibiotic (Betamox 120mg/kg) and analgesia (methadone hydrochloride 10mg/kg). Mice were closely monitored for 4 days after surgery and further analgesia was given daily for 1-2 days during recovery of the animal. Imaging and behavioural training were not started until at least one week after surgery.

### Immunohistochemistry and *ex vivo* imaging

Brains were fixed by transcardial perfusion with 4% paraformaldehyde in phosphate buffer 0.1 M, followed by 24h of postfixation in the same solution at 4°C. The whole brains were incubated successively in 15% and 30% sucrose in phosphate-buffered saline (PBS) at 4°C for 2 and 12h respectively. Brains were sectioned tangentially to the surface of visual cortex at 80µm thickness on a microtome (Leica). Slides were washed and permeabilized with 0.4% Triton X-100 in PBS for 4 × 15 minutes and then incubated with blocking buffer (0.3% Triton X-100 + 5% BSA + 10% Normal Donkey Serum and 10% Normal Goat Serum in PBS) for 3h at room temperature. Primary antibodies were incubated with blocking buffer (0.3% Triton X-100 + 1% BSA + 5% Normal Donkey Serum and 5% Normal Goat Serum in PBS) overnight at 4°C. The next day, slides were washed and incubated for 2h with secondary antibodies, then mounted in DABCO-PVA (2.5% DABCO, 10% polyvinyl alcohol (Sigma; Type II), 5% glycerol and 25 mM Tris buffer at pH 8.7). The slides were imaged with a confocal microscope (Zeiss LSM 800), and confocal z-stacks were compared with the previously acquired *in vivo* imaging planes and z-stacks of the recording sites. We determined the approximate location of the injection site using GCaMP7f fluorescence and then used blood vessel patterns and cellular morphology to identify the imaging site. We matched at least three points in the confocal z-stack to points in the *in vivo* imaging plane to obtain a three-dimensional transformation matrix that was applied to the entire confocal z-stack. Cells were then manually identified and assigned to cell classes based on immunostaining. Primary antibodies and dilutions used: Rat anti-Somatostatin, 1:200 (Millipore MAB354); Mouse anti-Parvalbumin, 1:5000, (Swant PV235); Rabbit anti-Vasoactive Intestinal Peptide (VIP), 1:500 (Immunostar #20077). Secondary antibodies and dilutions used: Goat anti-Rat Alexa 647, 1:500 (Thermo Fisher #A21247); Donkey anti-Mouse Dylight 405, 1:500 (Jackson Immunoresearch #715-475-150); Goat anti-Rabbit Alexa 594, 1:500 (Thermo Fisher #A11012).

### Two-photon calcium imaging

Two-photon imaging was performed using a custom-built resonant scanning two-photon microscope (Cosys) and a Chameleon Vision S laser (Coherent) at 930nm using a 16X, .8NA objective (Nikon). Images were acquired using a 12 KHz resonant scanner (Cambridge Technology) and an FPGA module (PXIe-7965R FlexRIO, National Instruments). Multi-plane imaging was performed using a piezoelectric objective scanner (Physik Instrumente). All recordings were made of neurons in L2/3 (generally 150-250um below the surface). Each imaging volume consisted of 6 planes, 20µm apart, approximately 450×450um, 512×512 pixels in size. Images were captured at an effective framerate of 6.3 Hz per volume. At the beginning of each session anatomical landmarks were used to find and record from the same imaging site as on previous days. Mice which were found to have bone regrowth under the window, poor viral expression or many brightly labelled cells with nuclear GCaMP7f expression were excluded from the study.

The coarse receptive field position of each imaging site was determined on the first imaging day to ensure the visual stimuli were approximately centred in the receptive field. The monitor in front of the contralateral eye (covering ∼100 × 60 degrees of visual space) was divided into a 4 × 3 grid and stimuli alternating between black and white at 2Hz were presented at each grid position on a grey background in randomized order (10 repetitions). Stimuli were generated using Psychtoolbox-3 in MATLAB. At the end of all *in vivo* imaging data collection, a high-quality image stack of all recording sites was acquired under anaesthesia to aid subsequent registration with immunohistochemically labelled brain slices.

### Behavioural training

The equipment and method used for behavioural training was similar to previous studies ^5,49^. Mice were trained first on a visual discrimination go-no go task followed by the full odour-visual attention switching task described below. Mice were food restricted to maintain at least 85% of their free-feeding body weight (typically 85-90%, 2-3g of standard food pellets per animal per day) but had free access to water. A 10% solution of soy milk powder (SMA Wysoy) was used as reward during the task and delivered through a spout positioned near the snout of the mouse. Licks to this spout were detected through a piezo disc sensor and reward was released by opening a pinch valve (NResearch), both controlled by custom electronics. The visual stimuli were presented on two luminance-corrected monitors (luminance meter LS-100, Konica Minolta) positioned at 45° angles and 25cm distance relative to the mouse. Visual stimuli were generated using psychtoolbox-3 and all behavioural tasks were controlled using custom scripts written in MATLAB and with a Teensy microcontroller board.

Mice were first habituated to handling and gentle restraint over two to three days and were then head-fixed and trained to run on a polystyrene cylinder (20cm diameter) for a further one to four days. Mice were free to run on the polystyrene cylinder during all awake recordings and their running speed on this cylinder was measured using an incremental rotary encoder (Kübler).

Once mice were running reliably on the wheel, they performed one closed-loop visual discrimination behavioural session during which the movement of the mouse on the wheel controlled the movement of visual gratings on the screen. After this closed-loop session all subsequent sessions were with fixed spatial and temporal frequency of the drifting gratings, and mice were trained to run for sustained periods of time to initiate trials - at least 2.8s with an added random duration drawn from an exponential distribution (mean 0.4s). The stimuli used for visual discrimination were two sinusoidal gratings drifting in the opposite direction to the direction of running, with a fixed spatial and temporal frequency of 0.1 cycles per degree and 2Hz respectively. Unless otherwise specified the rewarded and unrewarded gratings were oriented +/-15° relative to vertical, symmetrically on both screens. The stimulus presented on a given trial was selected at random.

The mouse could trigger the release of a drop of soya milk when the rewarded grating was displayed by licking the reward spout during the ‘reward period’. The reward period started 1.5s (with added random duration, mean 0.2s) after the rewarded visual stimulus onset and lasted until the offset of the stimulus 1s later. If the mouse licked during the ‘reward period’ the trial was recorded as a ‘hit’, if the mouse did not lick it was recorded as a ‘miss’ and a drop of soya milk was dispensed automatically shortly before the disappearance of the visual stimulus. Miss trials were rare after the first few days of training. A lick at any time when the unrewarded grating was displayed was recorded as a ‘false alarm’ and the mouse was punished with a 4s time-out period, during which the unrewarded grating persisted on the screen and any more licks reset the time-out duration. Ignoring the unrewarded visual stimulus by not licking was recorded as a ‘correct rejection’. In the initial stages of training, to discourage incorrect licking the probability of unrewarded trials was sometimes temporarily increased from 0.5 to 0.7. Mice typically learned the visual discrimination task in 5-10 days, with the threshold for learning defined as three consecutive days of discrimination with a behavioural d-prime score of 2.0 or above. Behavioural d-prime was calculated as: d’ = Φ^-1^ (H) - Φ^-1^ (F), where Φ^-1^ is the normal inverse cumulative distribution function, H is the rate of hit trials, and F is the rate of false alarm trials.

After learning the visual discrimination task, mice were trained to perform odour discrimination. The odour discrimination task was identical in structure to the visual discrimination task except that instead of visual stimuli one of two odour stimuli were presented to the mouse via polyethylene tubing positioned above the snout of the mouse. A custom-built flow-dilution olfactometer calibrated with a mini PID (Aurora) delivered 10-20% saturated vapour concentration of two solutions, 10% soy milk (rewarded odour) and 10% soy milk with 0.1% p-Cymene mixture (unrewarded odour). Mice typically started accurately discriminating between the odours after 30-40 trials, after which they were trained to switch between blocks of the olfactory and visual discrimination task.

Mice typically learned to perform the attention switching task in a further 1-3 days. In the olfactory blocks, 70% of odour stimuli were preceded by one of the two visual gratings presented in the visual discrimination task. These irrelevant visual stimuli were displayed for a fixed duration of 1.8s, with an onset delay distribution identical to the visual block, and neither grating was rewarded or punished. Mice learned to accurately discriminate between the odours while ignoring, i.e., not licking in response to either irrelevant grating. One of the two odours followed the irrelevant visual grating offset with a delay of 1.5s, plus an added random duration drawn from an exponential distribution with mean 0.2s.

### Optogenetic manipulations

Expression of the tdTomato conjugated opsin (Chrimson or ArchT) was first verified in each imaging site through two-photon imaging at 1030nm excitation wavelength. Optogenetic light was delivered using a digitally triggered 637nm laser (OBIS 637nm LX, Coherent), through a 200µm diameter 0.39 NA optic fibre (Thorlabs) positioned above the cranial window. To allow for quasi-simultaneous two-photon imaging and optogenetic activation, the laser and stimulus monitors were blanked during the linear phase of the resonant scanner.

For each two-photon imaging site an optogenetic calibration session was performed. During these calibration sessions the screens were grey, and 8 light powers (including 0%) were applied in pseudo-random sequence to the imaging window for 1.5s with 5s intervals. The effective maximum output used in mice expressing ArchT was 9mW, and in mice expressing Chrimson was 3mW. The average activity of each ROI in the 1s before the optogenetic laser onset was subtracted from the light period and the resulting baseline corrected activity was used for the calibration plots. Based on the shape of these calibration curves, 2 powers were chosen for the behavioural task for each imaging site: the lowest power producing a saturated or just below saturated response, and a second power with approximately half of this effect.

### Direction tuning

To examine the effect of the optogenetic light on visual processing in mice passively viewing stimuli, two-photon imaging sessions were conducted while head-fixed mice (free to run on a polystyrene wheel) were shown sinusoidal visual gratings drifting in one of 8 directions separated by 45°, with a spatial frequency of 0.1 cycles per degree and temporal frequency of 2Hz. These visual stimuli were randomly interleaved and one of three laser powers (including 0%, the same as those used in the attention switching task) was selected with equal probability for each visual stimulus presentation. The visual gratings were presented for 2s and the optogenetic laser lasted from 100ms before the stimulus onset to 1.5s after the start of stimulus presentation. There was a 5s interval before the start of the next visual stimulus presentation.

Direction tuning curves were constructed for each cell using their mean responses to each direction after baseline correction. The activity of each neuron was ‘soft’ normalized ^51^ so that neurons with strong responses had approximately unity firing rate range (normalization factor = firing rate range + 0.2).

When aligning direction tuning curves across neurons to each neuron’s preferred direction, the preferred direction was determined by taking the greatest response after pooling all light power and no light conditions. This prevented artefactual results which would occur from selecting the maximum firing rate in one condition (e.g., no light) and comparing this to other conditions (e.g., optogenetic stimulation). An orientation selectivity index was calculated as

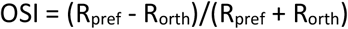

Where R_pref_ and R_orth_ are the average responses to the preferred and orthogonal directions respectively ^74^. Neurons were considered orientation selective if their OSI was greater than 0.33, such that the response at the preferred direction was twice as large as it’s response to the orthogonal direction.

### Optogenetic stimulation during attention-switching

Application of the optogenetic light during attention switching sessions was similar to the orientation mapping sessions. Optogenetic laser powers were interleaved and randomly selected on each trial from 1 of 3 powers (No laser, low power, high power) with equal probability. The optogenetic laser was on from 100ms before stimulus presentation to 1.5s after visual stimulus onset in Chrimson mice, or to the offset of the visual stimulus for ArchT mice. The optogenetic laser was delivered only during presentation of the visual stimuli in both the visual and odour blocks.

### Data analysis

Pre-processing of two-photon calcium imaging data was performed using the software Suite2p (https://github.com/MouseLand/suite2p) to correct for motion, detect regions of interest (ROIs) and extract the raw fluorescence time series of those ROIs, F(t). Each site yielded between 164 and 688 cells, median = 432 cells. We corrected the calcium traces for out of focus neuropil fluorescence using the neuropil masks identified by suite2p. For each frame we subtracted 0.7 * (neuropil - median neuropil fluorescence). Subsequent analysis unless otherwise specified was done with custom code in MATLAB and Python. Baseline fluorescence F_0_(t) was computed by smoothing F(t) (causal moving average of 0.75s) and taking the 2.5th percentile of the smoothed data. The change in fluorescence relative to baseline, ΔF/F, was computed by taking the difference between F and F_0_, and dividing by F_0_. Video recording (The Imaging Source) of the eye contralateral to the imaging site was performed during all sessions, and the time-points of saccades and blinks identified. Frames in which the mouse made a saccade or blinked were removed from further analysis. To identify VIP interneurons labelled with tdTomato *in vivo*, a brief dual channel recording of the imaging planes in red and green was taken before each imaging session at an excitation wavelength of 1020nm.

### Behavioural controls

To assess the proportion of neural activity which was attributable to overt behaviour recorded during our task, a linear model was fit using ridge regression to predict neural activity. The model was constructed by combining multiple sets of variables into a design matrix, to capture signal modulation by the following different task or behavioural events: 2 visual stimuli, 2 odour stimuli, reward delivery, licks, running speed, block type, and an interaction term for visual stimuli and block type. Each stimulus/event variable was structured to capture a time-varying event kernel. Variables therefore consisted of a vector of the relevant stimulus/event, and copies of this vector, each shifted in time by one frame for specific durations. For sensory stimuli, the time-shifted copies ranged up to 2s after the original. For motor events (running and licking) the time-shifted copies spanned the frames from 0.5s before until 2s after the original. The model was fit with 5-fold cross validation and the coefficient of determination (R^2^) was calculated based on the predictions of the model on held out data not used during training. We then assessed the predictive power of the behavioural model variables by comparing the R^2^ value for the full model to a model without the running and licking predictors, taking the proportion 1-(no behaviour model R^2^)/(full model R^2^).

To decode block type based on neural activity or running speed the neural decoding toolbox readout.info was used with the max_correlation_coefficient_CL classifier. To separate sessions according to their time-point of divergence in running speed the pre-stimulus baseline speed was subtracted and the first time-point found at which there was a significant difference in running speed between the attended and ignored rewarded visual stimulus trials using Wilcoxon rank sum tests at each time-point.

### Selectivity

We computed a selectivity index for individual ROIs as the difference between the mean response to each of the two gratings divided by the pooled standard deviation of that ROIs responses. Unless otherwise specified, all selectivity values presented here are from an analysis of the activity in the first 1s of visual stimulus presentation. To calculate ΔSelectivity we took the difference selectivity(attend) - selectivity(ignore). For cells that were negatively selective in the attend condition we multiplied the resultant values by –1, to ensure that cells that became more selective with attention had positive values. To test if an ROI was significantly selective within a certain time window, a two-sided Wilcoxon rank sum test was performed comparing the activity on trials for the two visual stimuli. ROIs were excluded from further analysis if they displayed selectivity in the period before visual stimuli were presented. To find ROIs that significantly changed their selectivity with attention, an ROI’s selectivity in the attend condition was compared to a distribution produced through bootstrapping using the data in the ignore condition (1000 repeats). If the attend selectivity was below or above the 2.5^th^ or 97.5^th^ percentiles respectively of the bootstrapped distribution, then the ROI was considered to have significantly changed its selectivity with attention. To avoid artefactual effects from selecting cells in one condition and testing in another, the test for significance was performed with no light and all light powers pooled.

After each behavioural block transition, transient periods of less accurate behaviour were discarded by identifying the trial in each block beyond which behaviour was stably accurate, that is, where mice displayed greater than 75% accuracy on both the go and no-go stimuli for the remainder of the block, and for the odour block, where mice licked in response to fewer than 25% of either of the irrelevant visual gratings for the remainder of the block. Light and no light trials were pooled to identify this point to avoid artefactual results.

### Orthogonality

Visualisation of the LDA transformation of neural activity was done using the Python library scikit-learn. All other analysis for testing orthogonality was done on axes found in MATLAB using the fitcdiscr function. The alignment of the axes best separating the optogenetic modulation and attentional modulation were found through the cosine similarity of the coefficients of two linear discriminant analysis models. One model separated visual stimulus trials in the odour block from visual stimulus trials in the visual block, both sets of trials without the optogenetic perturbation. The second model separated visual stimulus trials in the odour block without optogenetic perturbation from visual stimulus trials in the odour block with optogenetic perturbation.

The cosine similarity for the split data control was found using a similar approach. In the visual block, the rewarded and unrewarded visual stimulus trials were separated into two halves and the coefficients used to calculate cosine similarity were for two models separating the rewarded and unrewarded trials using non-overlapping halves of the data. This process was repeated 50 times for each session and the resulting values were averaged to produce one value for cosine similarity for split data for each session.

To ensure that we do not obtain orthogonal axes simply because they lie within a high-dimensional neural subspace, random axes were found using the method from ^51^. The neural covariance was estimated from each session and a Monte Carlo analysis used to sample pairs of random axes (10,000 samples) that were then used to calculate an expectation of cosine similarity based only on the dimensionality of the data.

### Noise correlations

To calculate noise correlation, the average stimulus evoked response across all trials of a particular type was taken for each cell and subtracted from each trial of the corresponding type. There were 14 trial types in total, rewarded and unrewarded odour stimuli, and rewarded and unrewarded visual stimuli in the visual and odour blocks, all at multiple laser powers. The Pearson correlation coefficient was then used to quantify the correlation between responses of pairs of cells for each trial type. Changes in noise correlations between different cell types with attention or optogenetic modulation were tested using a Wilcoxon signed-rank test on all sessions for which the post-hoc immunomatching had been successful (n = 4 mice).

### Circuit model

We modelled a circuit consisting of an excitatory population of layer 2/3 pyramidal cells (PYR), and three inhibitory populations, corresponding to PV, SOM, and VIP interneurons. The activity of the population *i* is described by its response *r*_*i*_, which evolves over time according to the following equation:

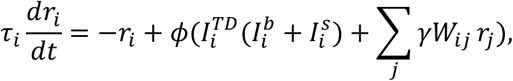

where *i, j* ∈ {*PYR, PV, SOM, VIP*} and

*τ*_*i*_ is the time constant of population *i*,

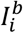 is the baseline input to population *i*,

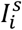 is the stimulus-dependent feedforward input to population *i*,

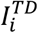 is the modulatory top-down input - the attentional modulation of population *i*, and

*Σ*_*j*_ *γW*_*ij*_ *r*_*j*_ is the recurrent input from the local circuit and *W*_*ij*_ is the effective synaptic weight from cell population *j* to *i*.

*γ* is a factor by which all weights are multiplied.

*ϕ*(*x*) is the activation function:

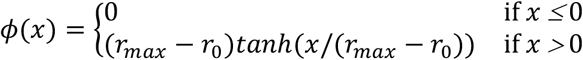

where *r*_0_ = 1.0 and *r*_*max*_ = 20.0 denote the baseline and maximum activity, respectively. PYR and PV populations received a stimulus-selective input current 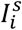 upon presentation of their preferred stimulus representing thalamic inputs. They received a fraction of this input current (0.5. *I*_*s*_) upon presentation of their non-preferred stimulus. Thus, the PYR and PV cell population activity corresponded to the average response of all cells with a given stimulus preference in the population. The SOM population also received a stimulus input current, which was the same for both presented stimuli. All populations received a constant baseline current input 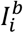. Each modulated population *i* (PYR, SOM, PV) received a multiplicative top-down modulation 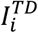 during attention (see Table 2). The VIP population did not receive top-down inputs in this model reflecting the lack of attentional modulation observed during the task (Supplementary Fig. 2).

To find parameter values consistent with the experimental observations, we used the python module sbi^56^, which implements neural inference algorithms for simulation-based inference. We used uniform prior distributions within the ranges shown in Table 1 from which parameter values were sampled. We ran circuit simulations to obtain training data consisting of the sampled parameter values and the output of the simulation for these values. This training data was then used to train a deep neural density estimator, provided by the sbi (v0.21.0) package (sequential neural posterior estimation – SNPE). The trained neural density estimator returns a posterior distribution *P*(*θ*|*x*) of the parameters *θ* given the desired model output *x*. To obtain the distribution of parameter values that are consistent with the desired model output (the experimental observations), we sampled parameter sets from the posterior. The observations we used to match the model were the PYR, PV, SST and VIP peak stimulus-evoked activity without the optogenetic light, the PYR, PV and SST peak stimulus-evoked activity with the optogenetic VIP activation, the increased PYR and PV activity to the preferred stimulus with attention, the decreased PYR and PV activity to the non-preferred stimulus with attention, and the increased PYR activity to both preferred and non-preferred stimuli with the VIP activation. For the final model, we took parameter values that had the maximum posterior probability.

**Table 1:** Sampling intervals for the parameters. The intervals from which parameter values were sampled for each parameter.

| Parameter | Interval | Parameter | Interval |
| --- | --- | --- | --- |
| $I_{PYR}^b, I_{PV}^b, I_{SOM}^b, I_{VIP}^b$ | [0,10] | $W_{EE}, W_{PE}, W_{SE}, W_{VE}, W_{PS}$ | [0, 1] |
| $I_{PYR}^s, I_{PV}^s, I_{SOM}^s, I_{VIP}^s$ | [0,10] | $\gamma$ | [1,5] |
| $I_{PYR}^{TD}, I_{PV}^{TD}, I_{SOM}^{TD}$ | [0,10] | | |

**Table 2:** Inputs to the cell types. The values for the baseline, stimulus, and top-down inputs to the populations PYR, PV, SOM, and VIP. Top-down modulation depends on the condition, which is either ignore or attend: {*x*_*ignore*_ *x*_*attend:*_}.

| Population | baseline $I_i^b$ | stimulus $I_i^s$ | top-down $I_i^{TD}$ |
| --- | --- | --- | --- |
| PYR | 6.0 | 3.3 | {1.0, 1.75} |
| PV | 4.9 | 0.1 | {1.0, 3.36} |
| SOM | 5.1 | 1.5 | {1.0, 1.76} |
| VIP | 5.3 | 0.0 | {1.0, 1.0} |

We inferred the following values of parameters for the baseline inputs 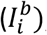, the feedforward inputs 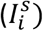, the modulatory top-down inputs 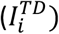.

For most values of connection strengths, we used the relative weights reported by ^17^. We multiplied all weights with a factor *γ*=1.485, which we inferred with sbi. Hence, the relative weights match the ones reported by ^17^, which were normalised such that *W*_*EP*_ was 1.00. We used simulation-based inference as described above to infer the parameter values of *W*_*EE*_, *W*_*PE*,_ *W*_*SE*_ and *W*_*VE*_ which were not reported by ^17^. Since the SOM to PV connection also plays an important disinhibitory role, we adjusted the value for *W*_*PS*_ to match the observations in the data. In our model the value of *W*_*PS*_ was larger compared to the value reported by ^17^ (0.5 vs. 0.3). Critically, with the smaller value, it was not possible to match the observations from the data.

The final connections between cell types were as follows:

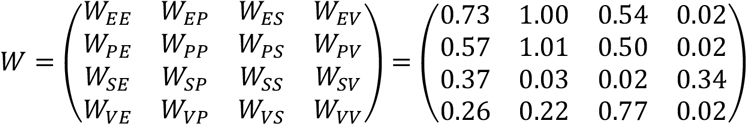

We simulated the network without stimulus input for 4s until the neural activity for each cell class reached steady state. Then we presented the stimulus (either preferred or non-preferred) for 1.5s. In the attend condition, we introduced multiplicative top-down modulation 2s after the start of the stimulation (2s before the stimulus). Optogenetic light stimulation happened at the same time as stimulus presentation. During light stimulation, we set the activity of the VIP population to a fixed value such that the normalized response was 2.0, in the case of VIP activation or 0 in the case of VIP inactivation. The simulation time step was 0.2ms. *τ*_*i*_ with *i* ∈ {*PYR, SOM, VIP, PV*} was 100ms. All responses were baseline-subtracted (baseline at 50ms before stimulus onset) and to ensure that the responses were comparable to fluorescence intensity measurements, the model responses were divided by a factor of 5.0.

To calculate the mean response for each cell type and each condition, we took the average of the response over the stimulus window. To calculate the selectivity of cell populations in the model, we subtracted the mean activity to the non-preferred stimulus 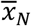 from the mean activity to the preferred stimulus 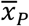 during the stimulus presentation:

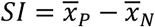

To study the impact of *W*_*PE*_ and *W*_*PS*_ on how much the VIP activation increased PV activity, we set either connection to 0.0 and compared the mean activity of the PV population in response to their preferred stimulus in the three conditions (control, without *W*_*PE*_, without *W*_*PS*_).

To visualize the parameter distributions of the top-down modulation used in the final model that is consistent with the desired model output we fixed the baseline and feedforward inputs, varied the parameters *W*_*EE*_, *W*_*PE*_,, *W*_*SE*,_*W*_*VE*_, *W*_*PS*_, *γ* and 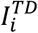, ran 10,000 simulations on which we trained the neural density estimator (SNPE), and then sampled 50,000 parameter sets from the estimated posterior distribution.

## Supplementary figures

**Supplementary Figure 1:**
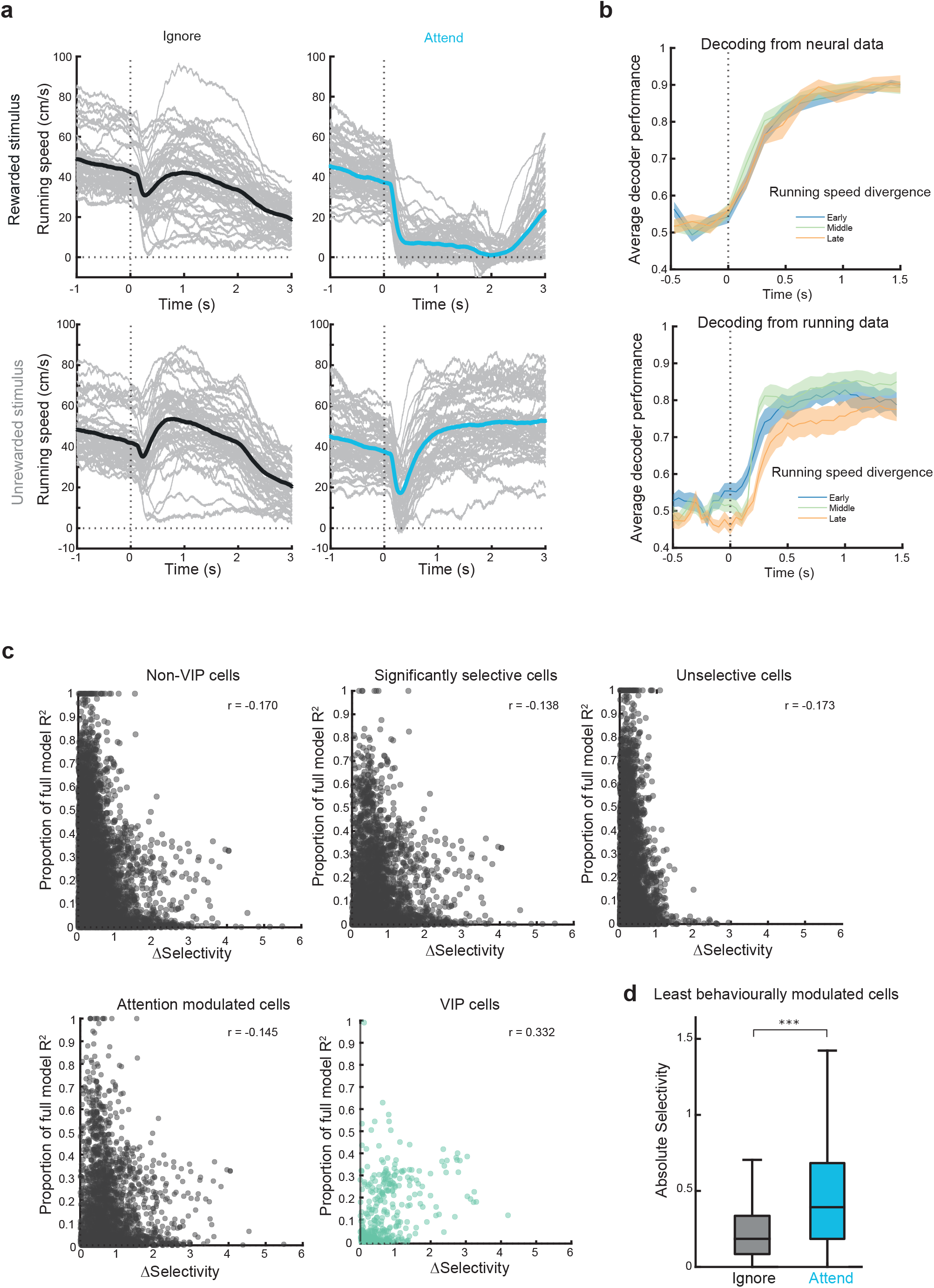
Changes in running and licking cannot account for the increase in stimulus selectivity with attention.**a**) Running speed aligned to rewarded and unrewarded visual stimuli onset (dashed line) - top and bottom respectively. Left - trials from the odour block, right - trials from the visual block. Grey traces are the individual session averages, coloured lines are the overall average. **b**) Top - performance of a maximum correlation coefficient classifier when decoding block type from neural activity evoked by the rewarded visual stimulus in 158ms bins aligned to the visual stimulus onset. The data was divided into three groups of sessions by the time of divergence of running speed between attend and ignore conditions. Early, middle and late represent 0-33^rd^, 34^th^-66^th^ and 67^th^-100^th^ percentiles. Bottom, same as top, but for a classifier using running speed instead of neural activity. While the decoder using running data clearly follows the sequence of running divergence, the decoder using neural data does equally well in all three running conditions, demonstrating that the running divergence does not contribute to the distinct activity levels in the attend and ignore conditions. **c**) Cells with stronger influence of running and licking do not account for attentional modulation. Relationship between absolute change in selectivity for the visual stimuli with attention (ΔSelectivity) and the reduction in R^2^ when removing running and licking behaviour information from a ridge regression model predicting the activity of each neuron: 1 - (no-behaviour model R^2^)/(full model R^2^). Larger values indicate a greater influence of running and licking. From top left to bottom right: All non-VIP cells, all significantly selective non-VIP cells, all non-VIP cells that were not significantly selective, all non-VIP cells that significantly increased their selectivity with attention, all VIP interneurons. None of the selections of non-VIP cells had a significant positive correlation, and in fact had small but significant negative correlations (Pearson correlation coefficients -0.17, -0.14, -0.17, -0.15 respectively, all ps < 10^−12^) demonstrating that selectivity modulation by attention cannot be accounted for by running and licking behaviours. VIP interneurons showed a significant positive correlation (Pearson correlation coefficient 0.33, p = 1.76×10^−10^), consistent with their well-established modulation by running. **d**) Average absolute stimulus selectivity during the ignore and attend conditions (n = 3086 cells, 15 mice, non-VIP neurons) when selecting only neurons whose activity is least influenced by running and licking behaviour (bottom 50% of the median change in model R^2^, Wilcoxon signed-rank test, p = 0).

**Supplementary Figure 2:**
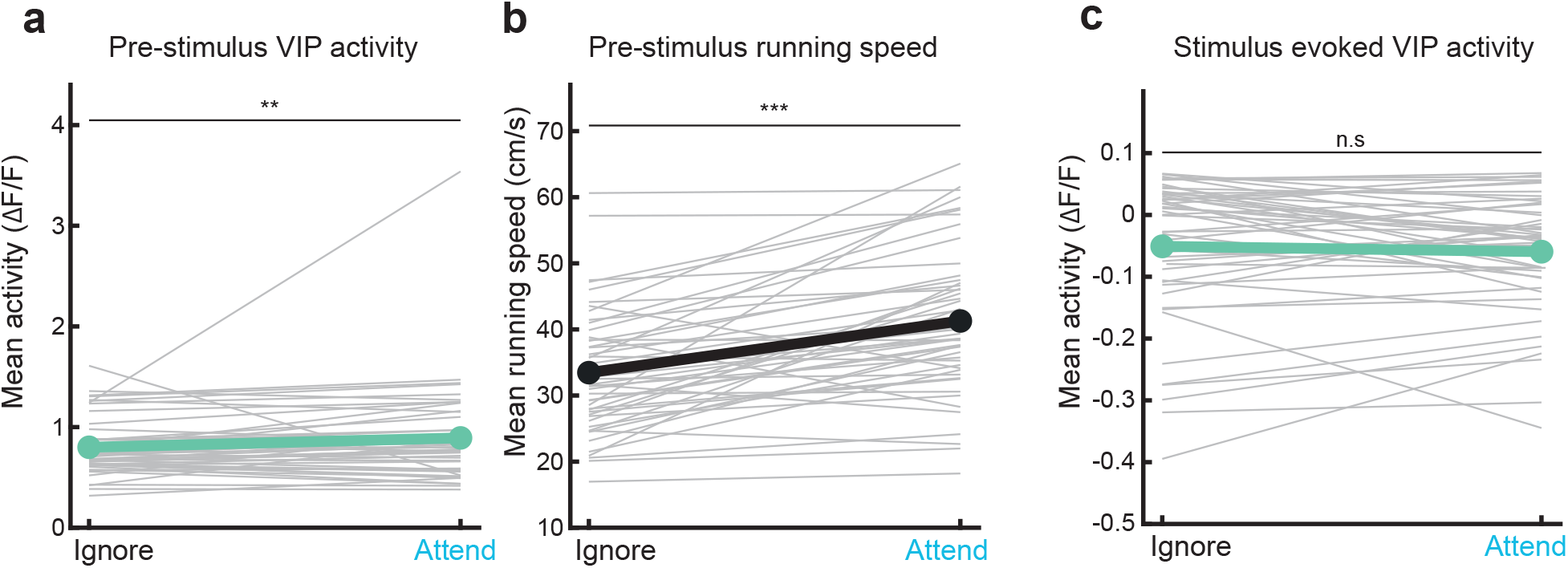
VIP interneuron activity in ignore and attend conditions.**a**) Average pre-stimulus VIP interneuron activity -0.1 to -1s relative to visual stimulus onset in the ignore and attend conditions of the task. Grey lines are the individual session averages, coloured lines are the overall average (**, p < 0.01, n = 50 sessions, 15 mice). **b**) Average running speed -0.1 to -1s relative to visual stimulus onset in the ignore and attend conditions of the task. Grey lines are the individual session averages, coloured lines are the overall average (***, p < 0.001, n = 50 sessions, 15 mice). **c**) Average stimulus evoked VIP interneuron activity 0-1s after visual stimulus onset (baseline subtracted) in the ignore and attend conditions of the task. Grey lines are the individual session averages, coloured lines are the overall average (n = 50 sessions, 15 mice, n.s. indicates not significant).

**Supplementary Figure 3:**
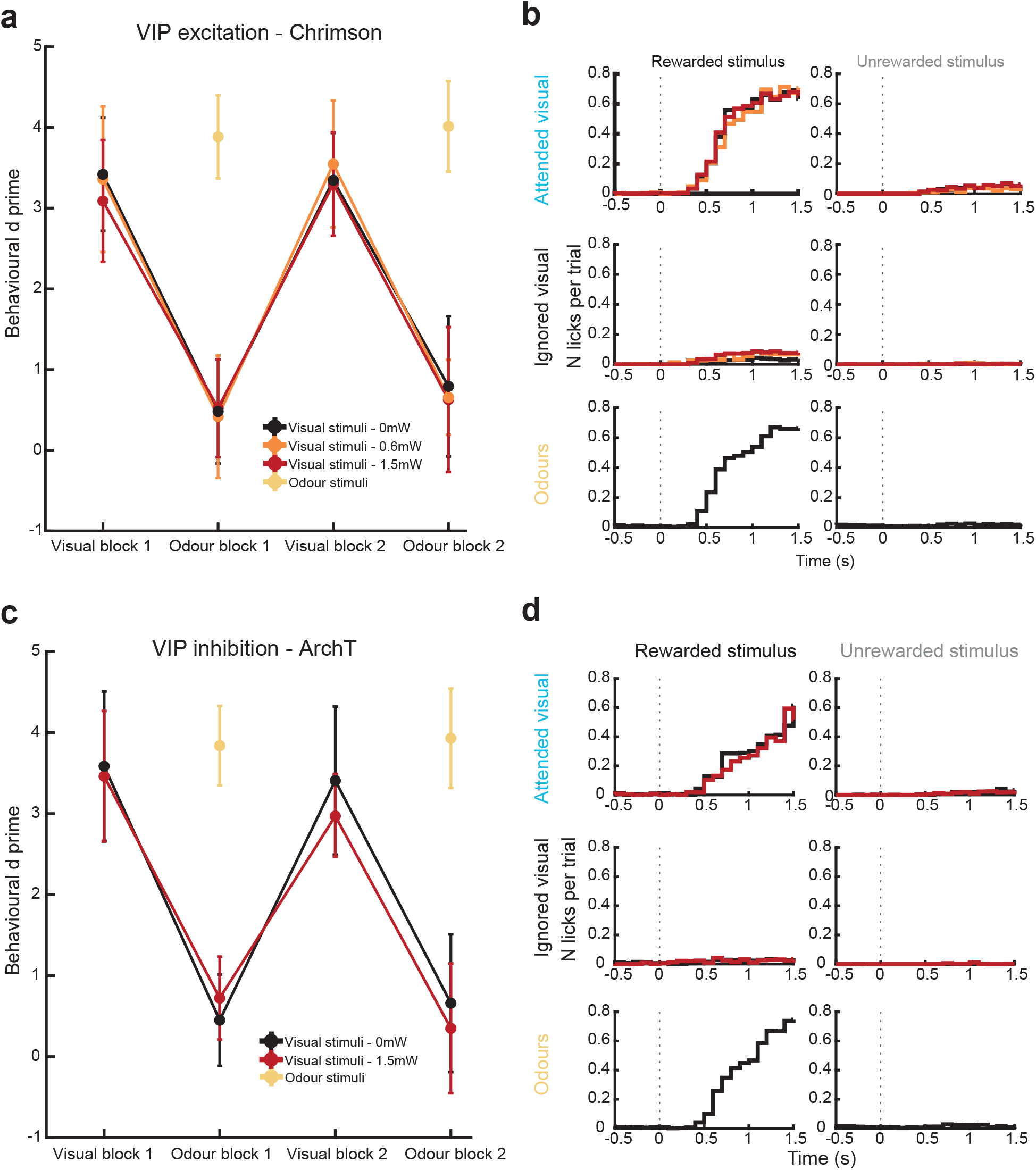
Behaviour is not affected by optogenetic manipulation of VIP interneurons in the imaging window.**a**) Behavioural d’ for visual and odour stimuli across the first 4 blocks of attention switching, shown separately for different optogenetic light powers, for mice expressing the excitatory opsin Chrimson in VIP interneurons. No changes were observed in the discrimination accuracy of visual or odour stimuli with optogenetic light. Wilcoxon signed-rank tests comparing visual stimulus no light and light trials: low power, visual block p = 0.801, odour block p = 0.461; high power, visual block p = 0.167, odour block p = 0.958. Error bars indicate SEM. **b**) Average number of licks per trial aligned to stimulus onset for each of 6 trial types, with trials divided according to optogenetic light power (same colour legend as a). Licks shown in response to the visual stimuli in the visual block (top), visual stimuli in the odour block (middle), and odour stimuli (bottom), split into the stimulus rewarded in the relevant block (left) and unrewarded stimulus (right). **c-d**) Same as in a-b, for mice expressing the inhibitory opsin ArchT in VIP interneurons. Wilcoxon signed-rank tests comparing behavioural d’ for visual stimulus no light and light trials: visual block p = 0.831, odour block p = 0.644.

**Supplementary Figure 4:**
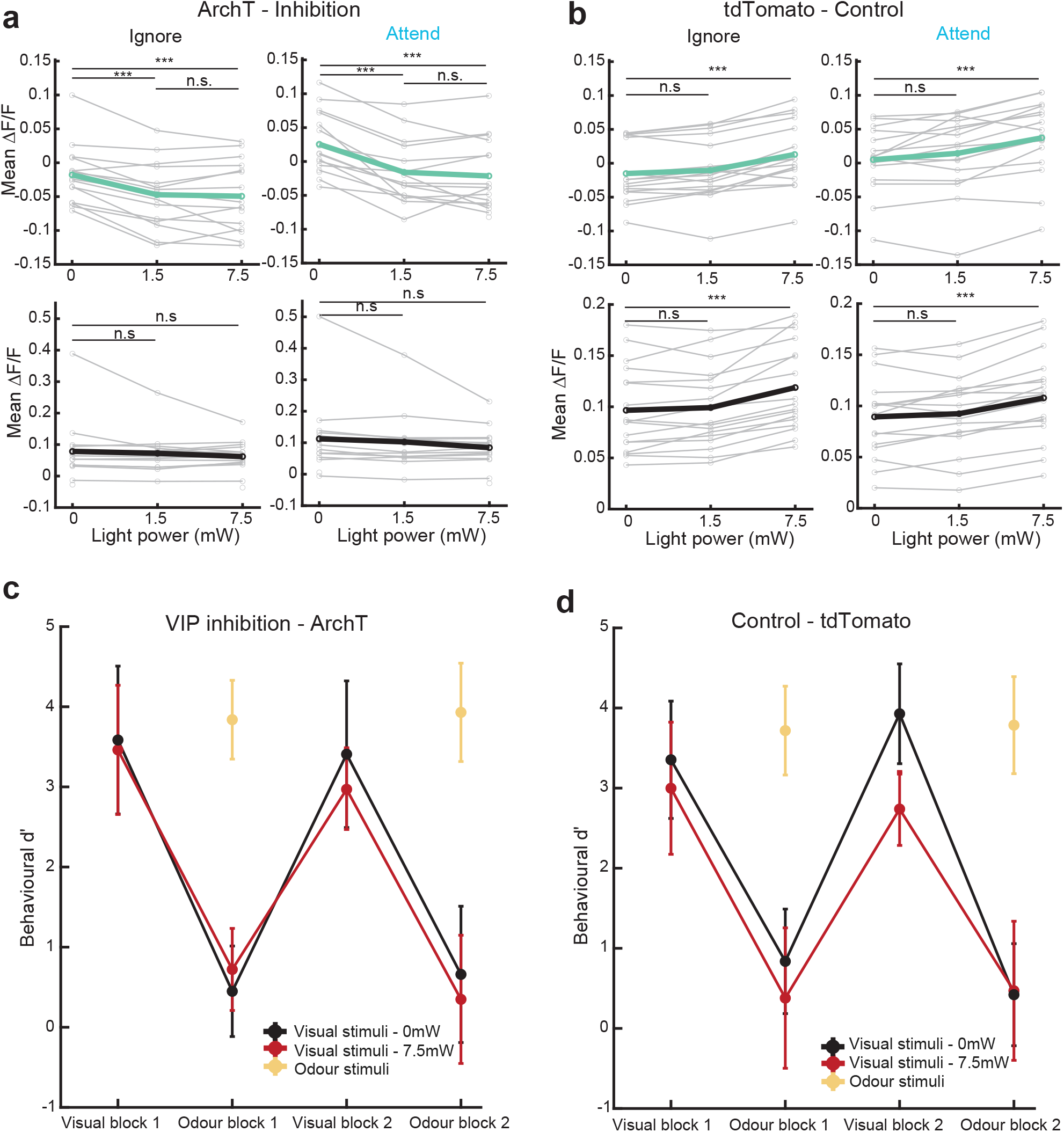
Increasing light power saturates VIP inhibition and introduces light-induced artefacts**a**) Average optogenetically evoked activity (baseline subtracted) with increasing light power for mice expressing ArchT in VIP interneurons. Top, all VIP interneurons, bottom, all non-VIP cells. Responses shown when ignoring the visual stimuli (left) or attending the visual stimuli (right). Wilcoxon signed-rank test between light activity and non-light activity, ***, p< 0.001, n.s. indicates not significant, n = 16 sessions, 5 mice. Gray lines indicate individual session averages, coloured lines indicate overall average. **b**) Same as in a, for control mice expressing only tdTomato in VIP interneurons (n = 17 sessions, 3 mice). **c**) Behavioural d’ for visual and odour stimuli across the first 4 blocks of attention switching, shown separately for different optogenetic light powers, for mice expressing ArchT in VIP interneurons. Wilcoxon signed-rank tests comparing visual stimulus no light and light trials: visual block p = 0.374, odour block p = 0.844. **d**) Same as c, for control mice expressing only tdTomato in VIP interneurons. Wilcoxon signed-rank tests comparing visual stimulus no light and light trials: visual block p = 5.07×10^−5^, odour block p = 0.132.

**Supplementary Figure 5:**
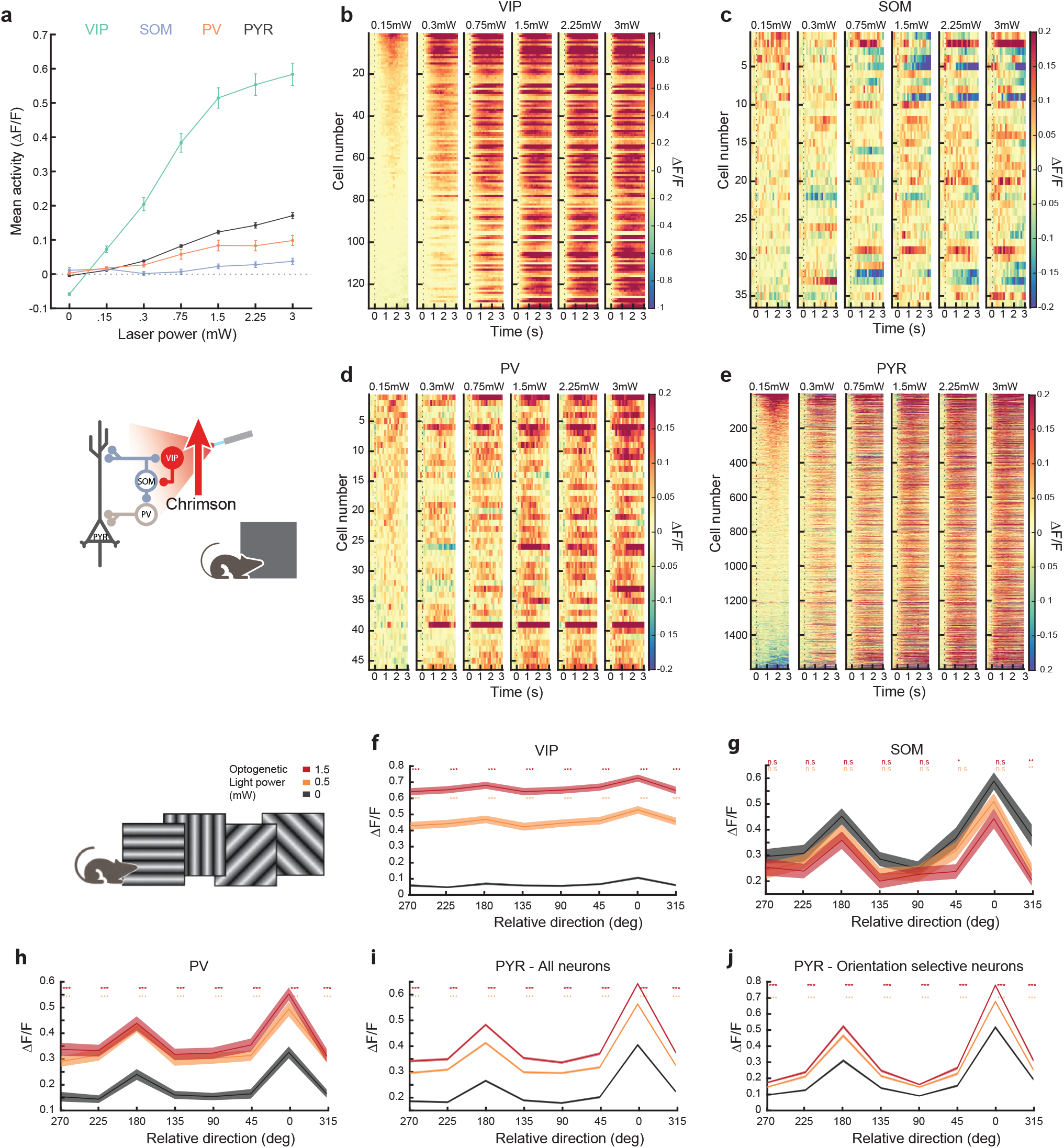
VIP photoactivation leads to changes in activity of multiple cell classes during passive grey screen viewing and presentation of drifting oriented gratings **a)** Average activity (0-1.5s) for all VIP, SOM, PV and PYR cells aligned to optogenetic light onset at each of 7 light powers. **b**) Average difference in activity between optogenetic light and no light trials for each recorded VIP interneuron at each of 6 optogenetic light powers, increasing left to right, n= 4 mice. Responses are baseline subtracted and aligned to the onset of the optogenetic light (dashed line). Cells sorted according to their mean activity in 0.5-1.5s in the no light condition. **c-e**) Same as b, for SOM, PV and PYR cells. **f**) Stimulus evoked normalised activity in response to different oriented gratings, averaged across all VIP interneurons (n = 85 cells, 3 mice) aligned to their preferred orientation. *, p<0.05, **, p<0.01, ***, p<0.001 Wilcoxon signed-rank test for photoactivation compared to non-photoactivation conditions at each orientation, corrected for multiple comparisons. Linear regression, low power, slope = 1.691, intercept = 0.347. High power, slope = 1.489, intercept = 0.567. **g**) Same as a, for SOM cells (n = 30 cells, 3 mice). Linear regression, slope = 0.858, intercept = -0.009. High power, slope = 0.690, intercept = 0.019. **h**) Same as a, for PV cells (n = 40 cells, 3 mice). Linear regression, slope = 1.158, intercept = 0.125. High power, slope = 1.304, intercept = 0.125. **i**) Same as a, for all PYR cells (n = 1567 cells, 3 mice). Linear regression, slope = 1.214, intercept = 0.074. High power, slope = 1.375, intercept = 0.092. **j**) Same as a, for orientation selective PYR cells (n = 472 cells, 3 mice). Linear regression, slope = 1.270, intercept = 0.034. High power, slope = 1.452, intercept = 0.042.

**Supplementary Figure 6:**
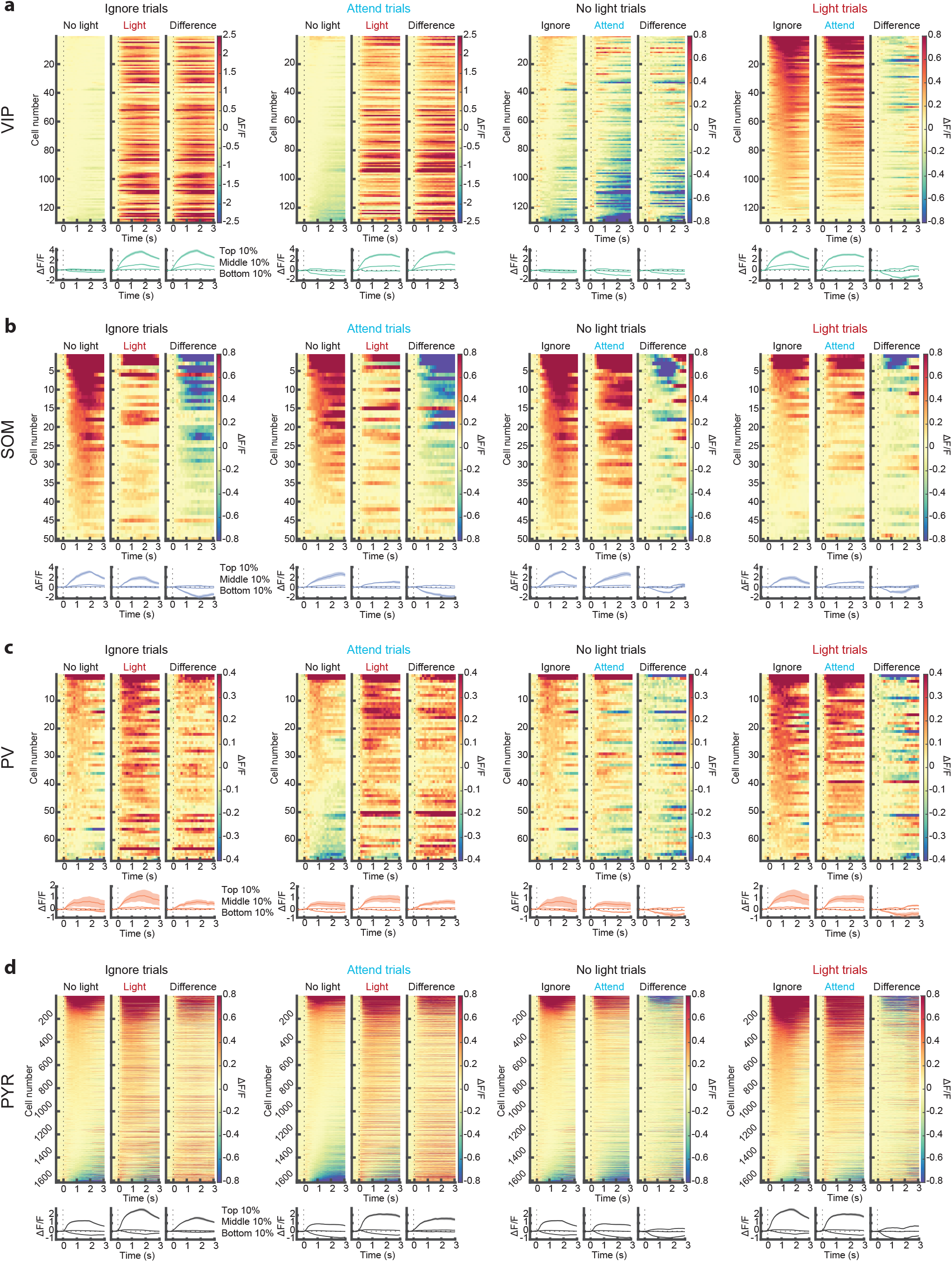
Changes in stimulus-evoked activity with attention and VIP photoactivation for multiple cell classes.**a**) Top, average visual stimulus-evoked responses for all VIP interneurons (n = 130 cells, 4 mice, baseline subtracted responses, average response to the unrewarded visual stimulus during the task) in two different conditions of no VIP photoactivation (No Light) and VIP photoactivation (Light) as indicated, and the difference of each pair of conditions. Cells are sorted according to their average activity (0-1.5s from stimulus onset) in the leftmost condition of each triplet. Bottom, average responses of cells from the top, middle and bottom 10^th^ percentiles of activity for each condition. Shaded areas indicate SEM. **b-d**) Same as a, for SOM cells (n = 50 cells), PV cells (n = 67 cells), and PYR cells (n = 1616 cells).

**Supplementary Table 1:** Two-way ANOVA results for the effects of attention and VIP activation on mean neural responses.

| Cell group | Visual stimulus | Attention effect | Light effect | Interaction |
| --- | --- | --- | --- | --- |
| Negatively selective | Preferred | p=0.133 | p=0.011 | F(2, 86) = 0.072, p=0.931 |
|  | Non-preferred | p=1.31x10<sup>-5</sup> | p=1.70x10<sup>-5</sup> | F(2, 86) = 0.376, p=0.688 |
| Positively selective | Preferred | p=0.316 | p= 0.005 | F(2, 86) = 0.016, p=0.984 |
|  | Non-preferred | p=0.080 | p=4.65x10<sup>-4</sup> | F(2, 86) = 0.017, p=0.983 |

**Supplementary Table 2:** Two-way ANOVA results for the effects of attention and VIP activation on neural stimulus selectivity.

| Cell group | Attention effect | Light effect | Interaction |
| --- | --- | --- | --- |
| All non-VIP | p=4.50x10<sup>-4</sup> | p=0.060 | F(1,64) = 0.955, p=0.332 |
| Unselective | p=0.002 | p=0.436 | F(1,64) = 1.200, p=0.278 |
| Significantly selective | p=0.012 | p=0.050 | F(1,64) = 0.512, p=0.477 |
| Increasing with attention | p=2.44x10<sup>-11</sup> | p=0.457 | F(1,64) = 2.095, p=0.153 |
| VIP cells | p=5.84x10<sup>-4</sup> | p=0.070 | F(1,60) = 1.351, p=0.250 |

**Supplementary Table:** Error! No text of specified style in document.: Two-way ANOVA results for the effect of attention and VIP inhibition on mean neural responses.

| Cell group | Visual stimulus | Attention effect | Light effect | Interaction |
| --- | --- | --- | --- | --- |
| Negatively selective | Preferred | p=0.076 | p=0.520 | F(2, 88) = 0.141, p=0.868 |
|  | Non-preferred | p=0.001 | p=0.792 | F(2, 88) = 0.028, p=0.972 |
| Positively selective | Preferred | p=1.74x10<sup>-11</sup> | p=0.667 | F(2, 88) = 0.017, p=0.983 |
|  | Non-preferred | p=0.033 | p=0.846 | F(2, 88) = 0.126, p=0.882 |

**Supplementary Table 4:** Two-way ANOVA results for the effects of attention and VIP inhibition on neuronal stimulus selectivity.

| Cell group | Attention effect | Light effect | Interaction |
| --- | --- | --- | --- |
| All non-VIP | p=1.72x10<sup>-10</sup> | p=0.886 | F(2,84) = 0.313, p=0.732 |
| Unselective | $p=6.76 \times 10^{-8}$ | $p=0.657$ | $F(2,84) = 0.967, p=0.385$ |
| Significantly selective | $p=2.58 \times 10^{-9}$ | $p=0.992$ | $F(2,84) = 0.048, p=0.953$ |
| Increasing with attention | $p=1.13 \times 10^{-16}$ | $p=0.878$ | $F(2,84) = 0.465, p=0.630$ |
| VIP | $p=3.38 \times 10^{-12}$ | $p=0.877$ | $F(2,84) = 0.006, p=0.994$ |

**Supplementary Table 5:** Two-way ANOVA results for the effect of attention and optogenetic light on mean neural responses in control mice expressing no opsin.

| Cell group | Visual stimulus | Attention effect | Light effect | Interaction |
| --- | --- | --- | --- | --- |
| Negatively selective | Preferred | $p=0.501$ | $p=0.887$ | $F(2, 96) = 0.365, p=0.695$ |
| | Non-preferred | $p=1.29 \times 10^{-8}$ | $p=0.762$ | $F(2, 96) = 0.022, p=0.979$ |
| Positively selective | Preferred | $p=1.84 \times 10^{-8}$ | $p=0.833$ | $F(2, 96) = 0.027, p=0.973$ |
| | Non-preferred | $p=0.088$ | $p=0.337$ | $F(2, 96) = 0.538, p=0.586$ |

**Supplementary Table 6:** Two-way ANOVA results for the effects of attention and optogenetic light on neuronal stimulus selectivity, in control mice expressing no opsin.

| Cell group | Attention effect | Light effect | Interaction |
| --- | --- | --- | --- |
| All non-VIP | $p=2.66 \times 10^{-10}$ | $p=0.741$ | $F(2,96) = 0.470, p=0.626$ |
| Unselective | $p=7.74 \times 10^{-6}$ | $p=0.961$ | $F(2,96) = 0.154, p=0.858$ |
| Significantly selective | $p=3.32 \times 10^{-5}$ | $p=0.922$ | $F(2,96) = 0.116, p=0.890$ |
| Increasing with attention | $p=2.69 \times 10^{-17}$ | $p=0.364$ | $F(2,96) = 0.734, p=0.483$ |
| VIP | $p=1.10 \times 10^{-14}$ | $p=0.406$ | $F(2,96) = 1.369, p=0.259$ |

